# Environment symmetry drives a multidirectional code in rat retrosplenial cortex

**DOI:** 10.1101/2021.08.22.457261

**Authors:** Ningyu Zhang, Roddy M. Grieves, Kate J. Jeffery

## Abstract

A class of neurons showing bidirectional tuning in a two-compartment environment was recently discovered in dysgranular retrosplenial cortex (dRSC). We investigated here whether these neurons possess a more general environmental symmetry-encoding property, potentially useful in representing complex spatial structure. We report that directional tuning of dRSC neurons reflected environment symmetry in onefold, twofold and fourfold-symmetric environments: this was the case not just globally, but also locally within each sub-compartment. Thus, these cells use environmental cues to organize multiple directional tuning curves, which perhaps sometimes combine via interaction with classic head direction cells. A consequence is that both local and global environmental symmetry are simultaneously encoded even within local sub-compartments, which may be important for cognitive mapping of the space beyond immediate perceptual reach.

**One Sentence Summary:** Retrosplenial directionally tuned neurons encode global environment symmetry even in local sub-compartments.

## Main text

Self-localization and navigation require construction of a stable directional signal that orients the brain’s map of space so as to be in register with the actual environment. Retrosplenial cortex (RSC) is a brain region where internal and external orientation signals combine, as evidenced by the long-established existence of head direction cells (*1*) which are compass-like, directionally tuned neurons that fire according to the animal’s facing direction. Recent findings revealed that in addition to these “classic” head direction (HD) cells, in dysgranular RSC (dRSC) there are also landmark-sensitive “bidirectional cells” which were found to express a bipolar firing pattern if recorded in a two-compartment space having twofold rotational symmetry (*3*). This bidirectional pattern was surprising, having not been noted previously, and we hypothesized that it arose from experience in the unusual environment structure, and reflects a general propensity for the cells to detect, and acquire encoding of, environment symmetry. To test this, we recorded RSC neurons in environments having symmetries varying between onefold, twofold and fourfold. We found that a subpopulation of electrophysiologically and anatomically distinct directionally tuned neurons showed multidirectional tuning that accorded with the environment’s symmetry, not just globally, when considered across the whole environment, but also sometimes locally within each sub-compartment. This pattern persisted in the dark and was not due to occult egocentric boundary encoding. Global multidirectional encoding indicates tuning of the neurons to the local environment polarities (and hence is only multifold when all the compartments are considered together), while local encoding of global symmetry means that the global multifold symmetry experienced after exploration has been captured by the network and is re-expressed locally. This enables spatial encoding of the environment beyond immediate perceptual reach, which may be important in building a cognitive map of a multi-compartment space.

## Results

RSC neurons were recorded from 18 rats (Table S1) foraging in environments having onefold symmetry (1-box), twofold symmetry (2-box) or fourfold symmetry (4-box). Every cell was recorded in one of the two 1-boxes and in either the 2-box or the 4-box (**Fig.1**; Fig. S1). We first present the 2-box data, then the 4-box, and then finally the 1-box results which flanked the multifold box trials.

**Fig. 1.**
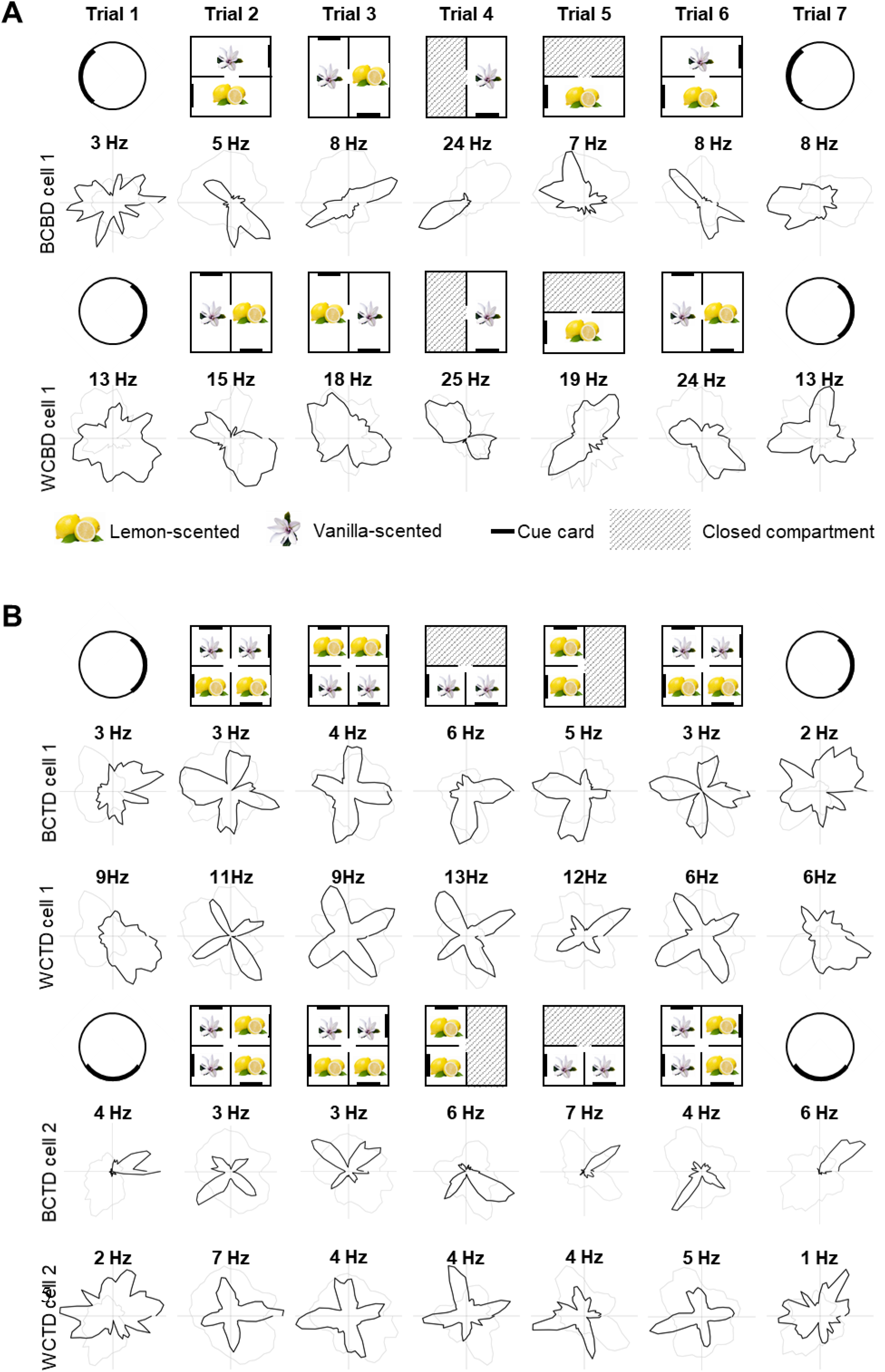
Directional tuning of BD-pattern cells in the 2-box experiment and TD-pattern cells in the 4-box experiment. (A) One between-compartment (BC) BD cell (top) and one within-compartment (WC) BD cell recorded in two different sessions. Polar plots show the standard seven-trial recording sessions: the rotation mark denotes the box orientation being different from that in the baseline (see Fig.S1 for full procedures). (B) Four TD cells: between-compartment (BC) TD cells and within-compartment (WC) TD cells recorded simultaneously from different tetrodes in one session (upper two rows) and another two co-recorded cells from a different session (lower two rows). Plots show cell activity in standard seven-trial recording sessions. Note that co-recorded cells have different PFDs within the same session. The decoupled HD tuning of co-recorded cells strongly supports that the firing symmetry is unlikely to be an artefact resulting from movement constraints of the animal during recording.

In the 2-box, we recorded 478 RSC cells from 14 rats (Table S1). As well as classic HD cells (27/478, 6%, **Fig. 2A-B**) and consistent with our previous work (*3*), we found directionally tuned cells with bidirectional tuning curves: a twofold-symmetric firing pattern with a 180º offset between two tuning curves (**Fig. 1A, Fig. 2C**; see Fig. S2 for more examples). This was quantified with a bidirectionality score and angle-doubled Rayleigh vector length (see **Methods**); 48/478 cells (10%) passed the criteria for bidirectionality. These cells displayed three main firing patterns. First, the so-called between-compartment bidirectional cells (BC-BD, n = 9/48, 19%, **Fig. 2C**; Fig. S1A) lost their bidirectionality in the local sub-compartments (**Fig. 2D**), as shown by the fact that BD scores were not significantly different from 0 (mean, lemon = 0.15, vanilla = −0.12, *p* > 0.3 in all cases); however they maintained unidirectional firing (median Rayleigh vector length: lemon = 0.22; vanilla = 0.23), as the within-compartment autocorrelations were single-peaked at zero and significantly different from a shuffle (KS test, lemon, D = 0.65, *p* <.001; vanilla, D = 0.59, *p* <.001). Thus, for these cells only the global pattern was bidirectional (**Fig. 2E**, cross-correlations significantly different from shuffle, KS test, D = 0.58, *p* <.001; peaked at 174°, circular v-test against 180°, V = 4.76, *p* = 0.012).

**Fig. 2.**
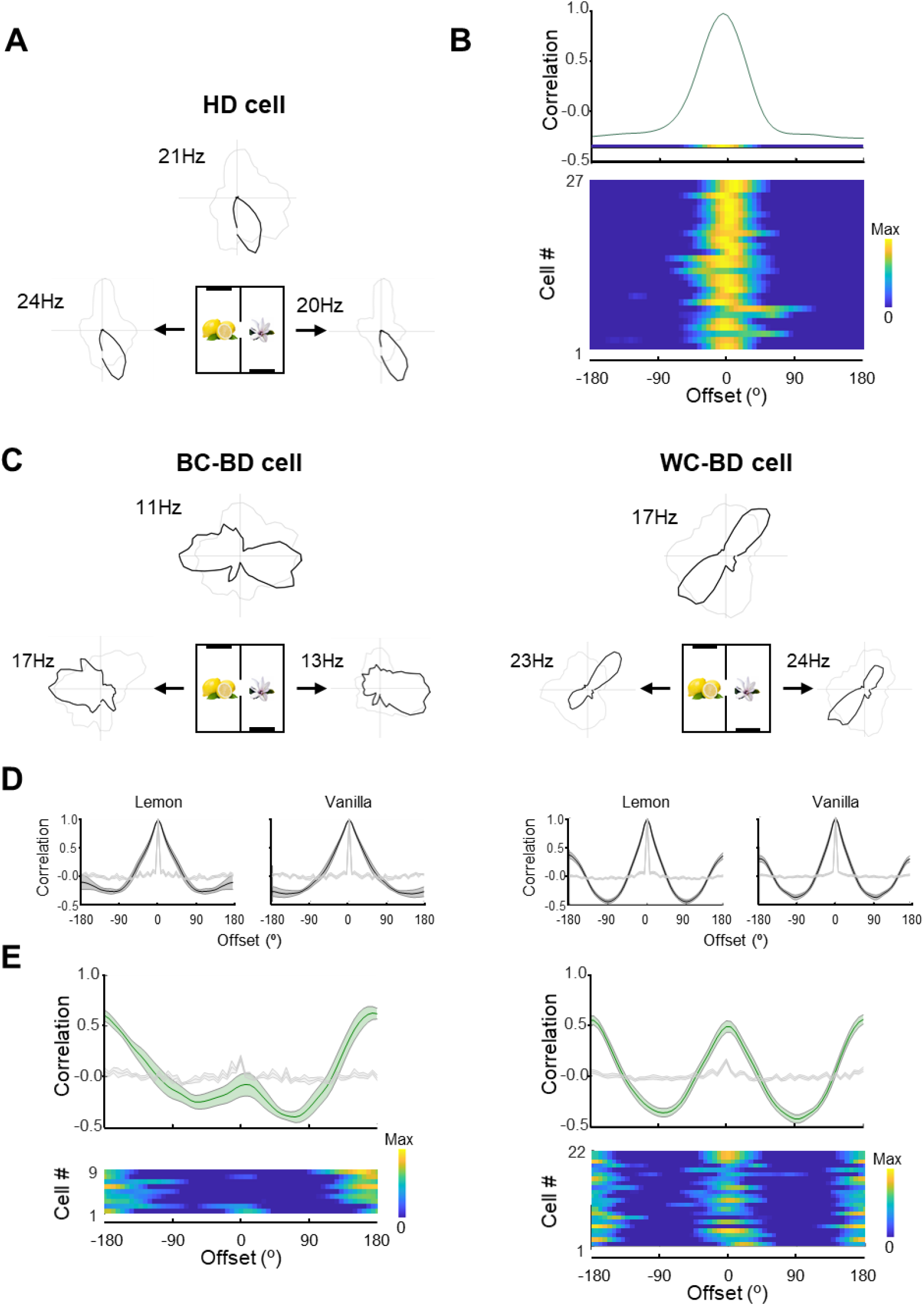
Multiple directional firing symmetries in the 2-box. (A-B) Classic HD cells. (A) Top polar plot: unipolar directional tuning curve for the whole apparatus (dark line) and directional behavioral sampling (pale line). Numbers show peak firing rates. Side plots: Tuning for each sub-compartment showing globally consistent direction. (B) Top: example tuning curve cross-correlation between sub-compartments, shown as a line and also as a heatplot. Bottom: heatplots of the cross-correlograms for individual cells, sorted by Rayleigh vector lengths; maximum at the top. (C) Global and compartment-specific polar plots of a between-compartment bidirectional cell (BC-BD; left) and a within-compartment cell (WC-BD; right). (D) Single-compartment autocorrelograms for the cells in (C) (mean +/- s.e.m.) showing a single peak for the BC-BD cell and double (second peak at 180 deg.) for the WC-BD cell. Shuffle control data are shown as pale gray. (E) Population cross-correlograms shown as line plot (top; shuffle controls as pale gray) and heatplots (bottom), again showing single peaks for the BC-BD cells and double for the WC-BD cells.

Second, the within-compartment bidirectional cells (WC-BD; n = 22/48, 46%, **Fig. 1A, Fig. 2C**, Fig. S2B), expressed a bidirectional pattern even within single sub-compartments (BD scores above shuffle control, mean lemon = 0.80; vanilla = 0.68; within-compartment autocorrelations significantly differed from a shuffle: KS test, lemon, D = 0.52, *p* <.001; vanilla, D = 0.52, *p* <.001; **Fig.2D**). Where the two within-compartment tuning curves were asymmetric, the direction of the large/small peaks rotated by ∼180° between the two sub-compartments (**Fig.2E**, between-compartment correlations significantly different from shuffle: KS test, D = 0.51, *p* <.001; mean ΔPFD = 188°, circular v-test against 180°: V = 6.54, *p* = 0.024). A third category of BD-pattern cells (17/48, 35%) fell below the statistical threshold for directionality in the sub-compartment analysis, and remained unclassified.

We next increased the level of environment symmetry from twofold to fourfold (the 4-box; Fig. S1). To rule out possible carry-over effects from the 2-box, we used four new rats, plus four rats that had previously had nine sessions in the 2-box (total rats, n = 8; cells, n = 660; Table S1). In the 4-box, in addition to HD cells (n = 75/660, 11%) we saw a fourfold-symmetric ‘four-leafed clover’ firing pattern, with a 90º offset between four directional tuning peaks (**Fig. 1B**). These also occurred in 3 out of 4 of the animals with previous 2-box experience (n cells = 15), suggesting pattern formation through learning of the new environment (Fig. S4). This fourfold directionality was quantified as a tetradirectional (TD) score, and 67/660 (10%) of cells met the criteria for tetradirectionality (**Methods**).

As with the BD cells, in single sub-compartments we identified three subgroups of TD-pattern cells: unidirectional, multidirectional and non-directional. First, the between-compartment TD cells (BC-TD; n = 9/67, 13%, **Fig. 1B, Fig. 3A**, Fig. S4A) expressed unidirectional but not tetradirectional tuning curves in individual sub-compartments (**Fig. 3B**). Unidirectionality was shown by a high Rayleigh vector score (median, vanilla1 = 0.27, vanilla2 = 0.22, lemon1 = 0.25, lemon2 = 0.32) and within-compartment autocorrelations significantly different from a shuffle in every sub-compartment (KS test, vanilla1, D = 0.67, *p* <.001; vanilla2, D = 0.66, *p* <.001; lemon1, D = 0.72, *p* <.001; lemon2, D = 0.52, *p* <.001). Loss of tetradirectionality was shown by within-compartment TD scores near zero (mean, vanilla1 = 0.00, vanilla2 = −0.02, lemon1 = −0.03, lemon2 = 0.03, *p* > 0.6 in all cases). The global preferred firing direction in each sub-compartment rotated by 90° in successive sub-compartments: cross-correlations of tuning curves from adjacent sub-compartment pairs differed from a shuffle (**Fig. 3C**, KS test, D = 0.68, *p* <.001) and peaked at ∼90° (circular v-test against 90°: V = 7.98, *p <*.001). These BC-TD cells also did not show a bidirectional pattern in single compartments, having BD scores not significantly different from zero (mean, vanilla1 = 0.07, t(8)= 0.51, *p* = 0.62; vanilla2 = 0.08, t(8)= 0.40, *p* = 0.70; lemon1 = 0.04, t(8)= 0.29, *p* = 0.78; lemon2 = −0.09, t(8)= −0.45, *p* = 0.67).

**Fig. 3.**
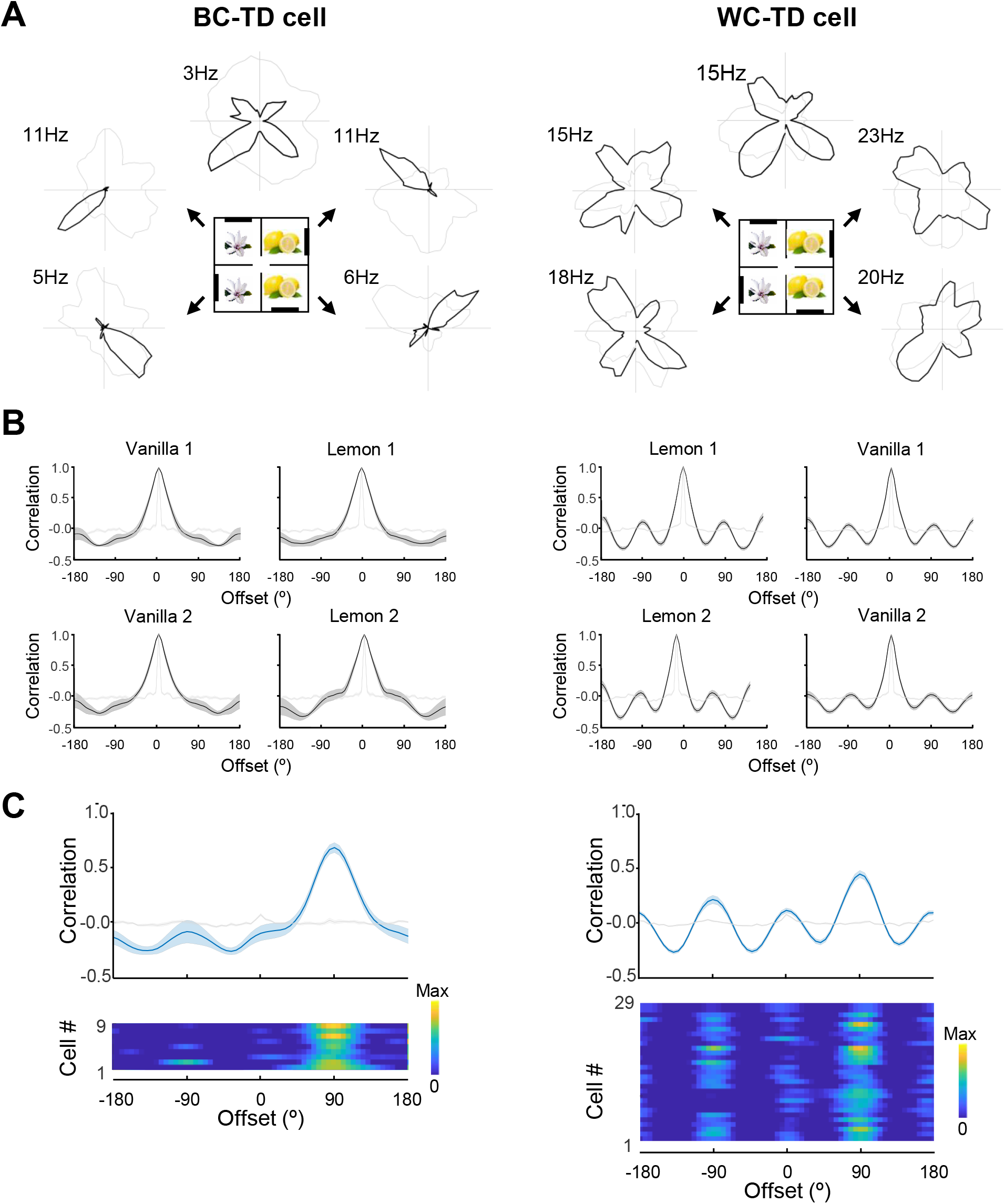
Multiple directional firing symmetries in the 4-box. (A) A between-compartment tetradirectional cell (BC-TD; left) and a within-compartment tetradirectional cell (WC-TD; right); illustrations as in Fig. 2. Note the occurrence of fourfold (90°) symmetries for the global pattern alone for the BC-TD cells, and for both global and local patterns in the WC-TD cells. (B) Rotational autocorrelograms of the tuning curves in each sub-compartment (mean ± s.e.m. = dark gray), compared with the shuffled data (light gray). The BC-TD cell has onefold symmetry (single peak) in each compartment while the WC-TD cell has fourfold symmetry (four peaks). (C) Population cross-correlograms (each compartment compared with the one clockwise adjacent): note peak at +90 degrees, reflecting rotation of the tuning curves to follow the rotation of the compartment layouts.

A second subgroup of TD-pattern cells, the within-compartment cells (WC-TD; n = 29/67, 43%, **Fig. 1B, Fig. 3A**, Fig. S4A) exhibited a multidirectional pattern (usually fourfold, though occasionally one peak was smaller) even in single sub-compartments. The within-compartment fourfold symmetry was quantified as follows. TD scores were significantly greater than the shuffle control: mean, vanilla1 = 0.41, t(28)= 6.03, *p* <.001; vanilla2 = 0.36, t(28)= 4.04, *p* <.001; lemon1 = 0.35, t(28)= 5.31, *p* <.001; lemon2 = 0.23, t(28)= 1.70, *p* = 0.05. Also, the coefficient of within-compartment autocorrelations was significantly different from shuffle: KS test, vanilla1, D = 0.49, *p* <.001; vanilla2, D = 0.47, *p* <.001; lemon1, D = 0.52, *p* <.001; lemon2, D = 0.50, *p* <.001; **Fig. 3B**. The tuning curve peaks varied in size, and the direction of the largest peak usually rotated 90º between successive sub-compartments. This was quantified using cross-correlograms of tuning curves from adjacent sub-compartment pairs, which significantly differed from a shuffle (**Fig. 3C**; KS test, D = 0.47, *p* <.001) and peaked at ∼90° (circular v-test against 90°: V = 13.69, *p* < .001). The third group (29/67, 43%) were not directional in individual sub-compartments and remained unclassified. We rarely saw cells with only two or three peaks in the whole 4-box. Moreover, we did not see any cells that were directionally stable across similarly-scented sub-compartments but that rotated between lemon and vanilla (i.e., there was no apparent hierarchy of sub-compartment encoding).

We also recorded the firing activity of TD-pattern cells in the dark in the 4-box (Fig. S5), finding that 75% maintained their four-way pattern in the first dark trial (n = 21/28), 64% in the second dark trial (n = 16/25), and 44% in both (n = 11/25). Together with previous findings of a preserved BD pattern in the dark (*3*), the results suggest that the multidirectional cells were not solely dependent on vision, and that multimodal sensory information could help maintain their directional tuning.

So far, we have shown that multidirectional cells are driven by a multifold environment symmetry, but how would they behave in environments with no repeating components? We thus next looked at the onefold-symmetry trials (the 1-boxes: the open arena and cylinder, Fig. S1), which flanked the multi-compartment trials. We found that while some cells lost directionality, a significant proportion became unidirectional: this was the case in both the open arena and the cue-controlled cylinder for cells that previously had been either BD or TD cells.

In the open arena, for cells that were BD in the 2-box (n = 32; **Fig. 4A**, Fig. S2), we found that overall, the cross-correlation between the two baselines had a tall central peak that significantly differed from the shuffle (KS test, D = 0.63, *p* <.001; **Fig. 4B**), indicating above-chance directionality at the population level. To quantify this at the level of individual cells, the Rayleigh vector length was compared with the cell’s shuffle in both baseline 1-box trials: 14/32 cells (were BD, 44%) had their Rayleigh vector lengths higher than its shuffle in both baseline trials (median, circular baseline 1 = 0.12, circular baseline 2 = 0.13), showing a broad unidirectional pattern; and 56% lost directional specificity. They did not show bidirectional pattern either: only 3/32 cells had BD scores higher than the shuffle.

**Fig. 4.**
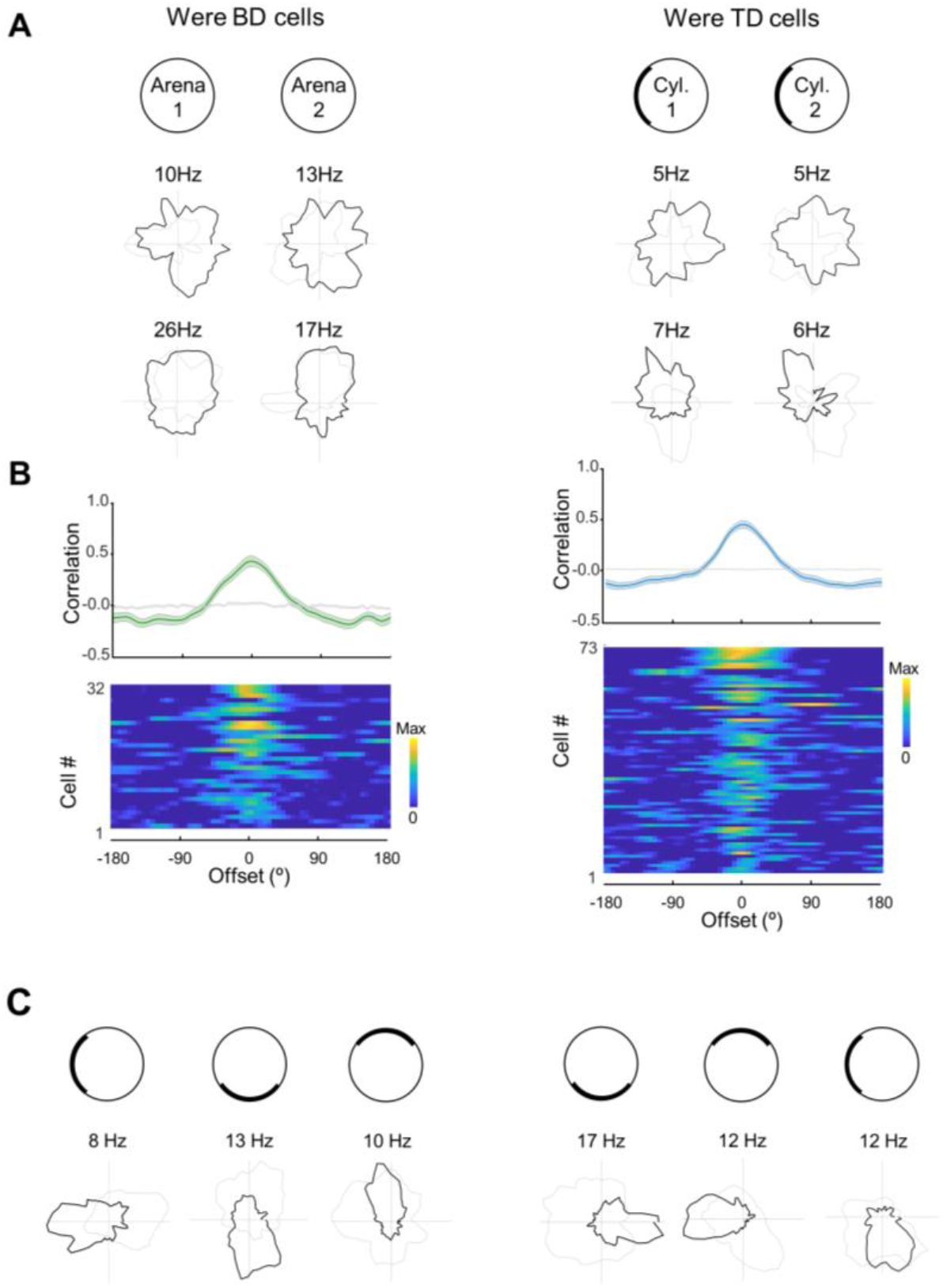
Variable uni-directional firing symmetries in the 1-box. (A) Two cells that had been BD cells in the 2-box (left) and two that were TD cells in the 4-box (right) showing varying directionality in the arena (left) vs. cylinder (right), either non-directional (upper cells in both examples) or unidirectional (lower cells). (B) Population data from former BD cells (left) and former TD cells (right) ordered top-to-bottom by Rayleigh vector; format as described in Fig. 2(E). prominent central stripe indicates preserved uni-directionality. (C) Two cells, a former BD-pattern cell (left) and former TD-pattern cell (right), which developed singular tuning curves in the 1-box that followed cue rotation.

We undertook a similar analysis for the cue-controlled cylinder 1-box trials (**Fig**. 1, **Fig. 4A**, Fig. S3, Fig. S6). Eight cells were formerly BD and 65 were formerly TD: we pooled these as “multidirectional”. As a population, the cells’ cross-correlations between the two 1-box baselines were significantly different from shuffle (KS test: n = 73 D = 0.67, *p* <.001; **Fig. 4B**). At the single-cell level, as before, the traditional Rayleigh vector length was compared with the cell’s shuffle in both baseline 1-box trials: 28/73 (38%) cells (26 were TD, 2 were BD) had their Rayleigh vector lengths higher than its shuffle in both baseline trials (median, circular baseline 1 = 0.17, circular baseline 2 = 0.18); and 45/73 (62%) cells lost the directional specificity. They did not show a multidirectional al pattern either in 1-box trials: only 1/8 had a BD score higher than shuffle, and 3/65 cells had their TD scores higher than the shuffle in both trials.

Some cells showed sensitivity to cue rotation (**Fig. 4C**) although this was inconsistent, perhaps due to familiarity of the rats with the room context. To assess whether this narrow onefold pattern is comparable with the unidirectional classic HD cells in the cylinder, we compared their peak firing rate, tuning width, and Rayleigh vector length, finding that these values were significantly different (Fig. S6): multidirectional cells showed a lower firing rate, lower Rayleigh vector score and broader tuning curves, suggesting that although they had become unidirectional, these cells were not the same as HD cells (see more extensive comparison below).

We then looked to see whether the multidirectional pattern we saw could be explained by occult tuning to the egocentric direction of boundaries, as has been previously suggested (*4*). Arguing against this possibility, we found that the allocentric directional tuning of retrosplenial cells was homogeneously distributed in the multifold environments, and therefore dissociated from the boundary directions which aligned along four distinct directions (Fig. S7A). Additionally, by applying established egocentric analyses (*4*) to the multidirectional cells, we found no selective egocentric boundary tuning in either the simple or complex environments: multidirectional cells generally showed multiple and uniformly dispersed egocentric firing fields (Fig. S7B-E).

We finally investigated whether multidirectional cells may be a physically separate neuronal sub-class from classic HD cells. First, and similar to previous results (*3*), more multidirectional cells were anatomically distributed in the dysgranular rather than granular RSC (Fig. S9; Table S1). Second, waveform analyses found that multidirectional cells in general had significantly higher peak amplitudes (ANOVA, *F*(2, 101.87) = 7.84, *p =* 0.001), longer peak-trough latencies (*F*(2, 100.98) = 159.73, *p*<.001) and longer latencies to the inter-spike-interval half-peak (*F*(2, 94.87) = 33.64, *p*<.001). Moreover, HD cells were significantly burstier than multidirectional cells (*F*(2, 140.61) = 24.81, *p*<.001). The results thus indicated an electrophysiological distinction between multidirectional cells and HD cells (Fig. S10; Table S2).

## Discussion

Our data show that a subpopulation of directionally tuned RSC neurons express an experience-dependent multidirectional code that captures the rotational symmetries of the extended environment. In addition to the well-known classic HD cells with globally unipolar, narrow directional tuning, some electrophysiologically distinct directionally tuned cells had firing patterns that varied according to the overall environmental symmetry, being one-directional in a single compartment but bidirectional in a twofold-symmetric environment and tetradirectional in a fourfold-symmetric environment. Furthermore, while some cells expressed these multidirectional patterns globally (being unidirectional in single compartments but multidirectional overall), others expressed them locally, expressing the global environment symmetry in local sub-compartments. These neurons thus provide an instantaneous readout of the global symmetry from within each of the sub-compartments.

Two questions arise from these findings: what generates these symmetries, and what, if anything, are they for? Clearly they are not intrinsic, because fourfold symmetric firing was not seen in the 2-box, and four- and twofold symmetry have not previously been observed in naïve animals in simple environments. However, we found that some cells that later became multidirectional expressed onefold directional tuning in the first 1-box baseline, suggesting a pre-configured responsiveness to local environment layout. Multidirectional firing was also found not to be due to egocentric boundary vector coding, and instead driven by allocentric environmental cues. The symmetries therefore evidently arise from experience of the different environment symmetries. However, they frequently persist in the onefold-symmetric sub-compartments afterwards, suggesting a learning process that was initially informed by the global exploration but then came to affect local activity.

Within-compartment expression of an across-compartment symmetry could potentially arise from plastic interactions with the HD system (*5*). Canonical HD cells use self-motion information to establish their globally stable directional orientation (*6*), while the non-canonical directional cells are sensitive to environment layout (*3*), developing multiple tuning curves if the environment has multiple identical but differently oriented sub-compartments. A consequence of this multidirectionality is that different populations of HD neurons become co-active with these neurons, depending on the global orientation of a given sub-compartment, and this co-activation could allow new HD inputs to form by Hebbian association, which then help to drive multidirectional activity.

Interestingly, the four tuning peaks in the fourfold-symmetric environment exceeds a proposed theoretical threshold (*5*) because previous computational modelling predicted that while the BD pattern would become tri-directional in a trifold-symmetric environment, the directional specificity should break down altogether with higher symmetries, due to the tuning curves overlapping (*5*). The fact that we saw fourfold symmetry suggests potential constraints on the parameters of the ring attractor that underlies the head direction signal (such as more narrowly dispersed interconnections).

What might this symmetry-influenced firing pattern be for? On the one hand it may be a byproduct of having explored a space with multiple similar-looking but differently-oriented compartments, forcing an unusual plasticity between the visual layout and the HD cells which confused the system. However there may be a deeper functional consequence, in which a global symmetry experienced when exploring multiple compartments can be re-expressed locally, for use in constructing an internal representation of a multi-compartment space. Such a construction, the so-called “cognitive map” of space, relies on recruitment of information beyond immediate perceptual reach. The ability of multidirectional RSC neurons to capture structural elements of the whole space (and their orientation) and then re-express them locally suggests that they may form part of such a mapping, perhaps as part of a compression process to more efficiently store complex structural information. In a more naturalistic environment, assuming that different cells are sensitive to different structural elements and that any repeating elements (such as corners, doors etc) would be irregularly distributed in directional space, the resultant population activity might look rather heterogeneous to an experimenter and yet contain a detailed plan of the global layout that could be use by other cognitive mapping neurons.

Overall, our results show that the subset of RSC directional neurons that become multidirectional in spaces with higher order rotational symmetries form a different subpopulation than classic head direction cells, and may play a role in mediating between the perception of local spatial features and construction of a cognitive map.

## Funding

This work was supported by a Wellcome Trust Investigator Award (WT103896AIA), a grant from the Biotechnology and Biological Sciences Research Council (BB/Joo9792/1) to K.J.J, and a China Scholarship Council Research Excellence Scholarship for PhD programme to N.Z. (201608000007).

## Author contributions

Ningyu Zhang: Conceptualization, Methodology, Software, Investigation, Formal analysis, Data curation, Visualization, Writing – Original Draft, Review & Editing; Roddy Grieves: Conceptualization, Software, Writing - Review & Editing; Kate Jeffery: Conceptualization, Methodology, Data curation, Supervision, Resources, Funding acquisition, Writing – Review & Editing.

## Competing interests

The authors declare the following competing interests: K.J.J is a non-shareholding director of Axona Ltd.

## Data and materials availability

A raw dataset can be requested from the authors. Summarized data and analysis code are documented in an online repository available upon publication.

## Supplementary Materials for

### Materials and Methods

#### Animals

A total of 18 adult male Lister Hooded rats (Charles River UK) were used in the study: 14 in the 2-box experiment, 8 in the 4-box experiment, with 4 in both; all also in one of the 1-boxes. They were individually housed in a temperature (22±2°C) and humidity-controlled (55±10%) holding room with a 12-hour light-dark cycle, with 1h of simulated dusk/dawn. A hanging hammock or nesting box was provided as enrichment. Animals received an implant surgery at a weight range between 350g-550g. Animals had free access to food and water during a seven-day recovery period after the surgery, then they were put on a mild food restriction diet (to 90% of free-feeding weight) throughout the experiment period. All procedures were carried out in compliance with the Animals (Scientific Procedures) Act, 1986, United Kingdom and the European Communities Council Directive of 24 November 1986 (86/609/EEC) legislation for use and maintenance of laboratory animals.

#### Implant surgery

Prior to surgery, each animal was subcutaneously injected with an analgesic (0.2ml Carprieve, Norbrook Laboratories, UK) mixed in a sterile saline solution (2ml, 0.9%) before being anesthetized and then stereotaxically implanted with tetrodes in either left or right RSC (target coordinates: AP −5.0-5.6mm, ML ±0.6-1mm, DV 0.5-0.7 mm, relative to Bregma). For half of the animals, the microdrive (Axona Ltd, St Albans, UK) was positioned at a angle of ∼10º dorsoventrally so as to sample both dysgranular and granular sub-regions. Animals received meloxicam for postoperative analgesia for 3 days.

#### Experimental apparatus

Recording enclosures of varying symmetry were used in the study, each placed in the center of a curtained-off cue-controlled region except for the circular arena, which was open to the room. Box schematics can be seen in Fig. S1. For the 2-box, we used the same apparatus as in the previous study (*3*) – two abutting 120 × 60cm rectangular boxes sharing a long wall that had a central doorway connecting the compartments. Each box was adorned with a single white cue card on one of the short walls, one at the “North” and one at the “South” end. This arrangement created a 180-degree rotational symmetry of the global space that was broken by scenting one compartment with lemon and one with vanilla odor. The 4-box (120 × 120 × 60 cm) contained four equal-sized square sub-compartments interconnected with a central 10 cm wide doorway. Each sub-compartment was polarized by two environmental features: the doorway in the innermost corner, and a white cue card (20 × 40 cm) mounted on the right-most facing wall as seen from the doorway. The spatial relationship between the cue card positions relative to the doorway remained the same for all four sub-compartments, forming a 90° rotational symmetry. At the start of each recording session, two adjoining sub-compartments were scented with lemon odor and the other two with vanilla odor made from bakery concentrates (Dr. Oetker, Bielefeld, Germany). The scents were intended to break the overall symmetry to enable HD cells to establish a stable orientation, and also to hierarchically cluster the sub-spaces in case there might be neurons showing a global directional tuning for a similar pair of subspaces. The floor and walls of the apparatus were covered with black vinyl sheets that were cleaned after each session. The multi-compartmented boxes were placed in the same experimental room, with floor-to-ceiling black curtains surrounding the apparatus to minimize distal cue influence.

For the onefold environments, two circular arenas (1-boxes) were used, located in two different experiment rooms. A bigger and low-walled circular arena (diameter = 100 cm, height 10 cm) was used with the 2-box only. It was raised 50 cm up from the floor and open to the distal room cues. The high-walled cylindrical arena (diameter = 80cm, height = 72 cm) was in different a cue-controlled room and used with all 4-box experiments and a few sessions in the 2-box experiment.

#### Electrophysiological recording procedures

Single-unit activity was recorded using a multichannel data acquisition system (Axona Ltd, St Albans, UK), monitored online with thresholds set by the experimenter. The microdrive was attached to a headstage, which was connected to recording cable leading to a pre-amplifier and digitizer (48 kHz). Single-unit signals were amplified 5,000-8,000 times and band-pass filtered between 300 and 7,000 Hz; local field potential (LFP) signals were amplified 1,000-2,000 times and lowpass filtered at 500 Hz. The head direction and position information of the animal was determined by tracking the positions of two LEDs (one brighter than the other) spaced 5 cm apart. Position and heading were monitored by an overhead camera, at a sampling rate of 50Hz. A radio positioned above the apparatus emitted constant white noise to mask directional auditory cues from beyond the curtains. During recordings, the environmental sound level was maintained over 70 dB to minimize any extra-maze auditory influence.

Before the start of the experiment, animals were acclimated to a separate screening room and were monitored daily for single-unit activity in circular environments. Tetrodes were lowered for 50-150 μm after every recording session or if no cell activity was detected. One standard recording session consisted of seven trials: the first and last trial in the 1-box and five multi-compartmented box trials in between:

‐ Trial 1 (1-box baseline): For high-walled cylinder trials, the cylinder was placed in the center of the room and oriented in multiples of 90° in camera-defined coordinates at the beginning of the trial. The aim of pseudo-random rotation in a cue-controlled room was to check whether the cell tuning follows the local cue rotation with the apparatus.
‐ Trial 2 (door-open box baseline): The apparatus was placed in the center of the room and oriented in multiples of 90° in camera-defined coordinates at the beginning of the trial. Animals were placed in a randomly chosen sub-compartment at the beginning of the trial.
‐ Trial 3 (door-open rotated box): As in trial 2 except that the apparatus was variably rotated by +/- 90° or 180°.
‐ Trial 4 (door-closed half box1): The central doorway was closed so that the animal was restricted to only half of the apparatus. The two lemon or two vanilla compartments were either adjacent or diagonal, to test for the possibility that there might be directional neurons that would be consistent across a two-compartment space if these were unified by a common odor (exposing a hierarchical encoding of sub-spaces). We found that contextual odors were not necessary for multidirectionality (see Fig. S2A).
‐ Trial 5 (door-closed half box 2): Same as in trial 4, but the animal was recorded in the other half of the apparatus.
‐ Trial 6 (door-open box baseline): As in trial 2, the door was open, and the apparatus was rotated back to its starting orientation.
‐ Trial 7 (1-box baseline): As in trial 1, the animal returned to the circular environment. All recording sessions included in the study consisted of at least the seven trials as described above. In addition, the following procedures were sometimes performed:
‐ Darkness trials: To test whether the firing pattern was primarily dependent on vision, in some 4-box recording sessions (n = 28), two darkness trials were added after the last door-open box baseline trial. The animal and cue cards were removed from the apparatus and the lighting sources in the room were switched off, then the animal was disorientated and reintroduced into the apparatus. The 4-box was randomly rotated before the second darkness trial.
‐ Rotated cylinder trials: In some recording sessions of the 2-box (n = 8) and 4-box experiments (n = 43), at least two extra cylinder trials were added following the last cylinder baseline trial to investigate whether the (uni)directionality derives from the visual environment. The rat was removed from the environment and mildly disoriented, the cylinder plus cue card were randomly rotated from the baseline orientation and then the rat reintroduced into the apparatus.

In all trials, animals were motivated to explore and sample all heading directions in the apparatus as much as possible while foraging for randomly scattered rice or coco-pops (Kellogs, UK). Between trials, the animal was gently removed from the apparatus, placed in a holding box inside the curtain to prevent its knowledge of any room cues or experimenter’s manipulation of the apparatus, and then mildly disoriented inside a custom-built rotating box before the next trial. Experiment room change happened between the trials in the 2-box/4-box and simple environments, and the animal was carried between the rooms in an opaque box to prevent perception of any distal cues.

#### Electrophysiological data analyses

##### 1. Cell identification

Single unit activity was analyzed offline using an automated spike sorting algorithm (Klustakwik v3.0)(*7*), packaged in cluster-cutting software with a graphical user interface (TINT, Axona Ltd, St Albans, UK). Cell cluster quality was assessed over a recording session that consists of multiple trials recorded from different environments. Before cluster-cutting, spike data collected from all trials within a session were compiled to one overall cluster space, so that the cluster center of mass remained constant across trials. Cluster quality was compared between uni- and multi-directional cells to check that cell isolation differences could not explain the findings. We did this by calculating three metrics: the isolation distance, L-ratio (both measures make use of the Mahalanobis Distance) (*8*) and refractory period contamination rate (*9*). The refractory period contamination rate was calculated as the number of spikes in the refractory period (2 ms duration) in proportion to the total number of spikes within a trial.

##### 2. Directionality analyses

‐ *HD polar plot:* For each trial, all spikes and head-direction samples were binned into 60 bins of 6° width, the number of spikes emitted in each bin divided by the total dwell time for that direction. A running average boxcar kernel (width = 5) was applied to smooth the raw tuning curve. The peak firing rate was defined as the maximal firing rate found in the smoothed tuning curve, and the angular bin of this maximum value was the cell’s preferred firing direction. For multidirectional cells the preferred firing direction was taken as the angle of the biggest peak. We tested PDFs from all directional cells for non-uniformity with a Rayleigh test (MATLAB function *circ_rtest* (*10*)).
‐ *BD/TD score:* The multidirectional scores reflected the level of spatial periodicity in the tuning curves and were calculated using a similar circular autocorrelation procedure as described previously (*3*). Each cell’s tuning curve was rotated against itself in steps of 6°, and the Pearson’s correlation coefficient re-calculated at each step. Periodicity in the tuning curve would result in a sinusoidal modulation of the circular auto-correlogram, and the number of peaks in this autocorrelogram would reflect the order of symmetry in the tuning curve. The bidirectional (BD) score tested the strength of twofold modulation and was defined as the difference between the mean correlation coefficient at the angle of the expected peak (±180°) and the average of the expected troughs (±90°). The tetrodirectional (TD) score tested the strength of fourfold modulation and was calculated similarly, but the expected peaks were at ±90° and ±180° and the expected troughs at ±45° and ±135°. Computation of the scores for the shuffled controls (see Shuffling section below) was performed in the same manner.
‐ *Directional specificity:* Normally, directional specificity was assessed using mean resultant vector length (Rayleigh vector length) (*11*). However, multi-fold directionality violates the unimodal assumption of the Rayleigh distribution: we thus performed the directionality analysis by angle-doubling (as used before (*3*)) in and angle-quadrupling procedures in which the heading direction of each spike (in radians) was doubled or quadrupled and wrapped at 2π. In this way, we converted multi-fold symmetric data into a unimodal distribution (*12*). Rayleigh vector length was then computed from the converted data. For the 1-box, Rayleigh vector length (without angle conversion) was computed for each cell and compared with the 95^th^ shuffle threshold in two 1-box baseline trials. To characterize the narrow unidirectional tuning pattern in the 1-box, the Rayleigh vector lengths of the cell should exceed its shuffle in three consecutive 1-box trials.
‐ *Shuffling:* A shuffling procedure was used to obtain control distributions and chance-level estimation in directionality and egocentric tuning analyses. In the HD analyses, for each permutation trial, the entire spike train of each cell was time-shifted by random intervals ranging from 20s to the trial duration (usually 600s for 2-box and circular 1-box, 960s for 4-box, and 480s for cylinder 1-box) minus 20s relative to the positional/directional data. Spike times shifted past the end of the trial were wrapped around to the beginning. Similarly in the egocentric analyses (see below), the random shifting interval was from 30s to the full length minus 30s relative to the positional data (the same parameters as used before (6)). One hundred permutations were performed for each trial of a given cell in the shuffling procedure. For each permutation, a shuffle HD tuning curve were constructed, and Rayleigh vector length, TD/BD scores, auto-correlograms, cross-correlograms, egocentric firing rate map and mean resultant length were determined on an individual basis. Thresholds were identified from the overall distribution of the measurements of the shuffled data (e.g., 95th percentile). At a population level, the population thresholds of Rayleigh vector length and multidirectional scores were the mean or the median 95th percentile threshold of all recorded cells in three door-open box trials. This is to obtain a robust threshold cutoff: the shuffle procedures were done for 478×100×3 times in the 2-box experiment and 660×100×3 times in the 4-box experiment. Note that the mean Rayleigh vector length was higher, and the median of multidirectional scores was higher, so more stringent thresholds were taken.
‐ *Selection criteria:* A classic head direction cell was defined as meeting the following criteria in both box-baseline trials: peak firing rate at least 1Hz and Rayleigh vector length greater than 0.26, the same threshold as used previously (*3*). For an overall classification of multidirectional cells, a BD-pattern cell was defined as meeting the following criteria: a). peak firing rate ≥ 1 Hz; AND b). BD score exceeds 95th percentile criterion of shuffle; AND c). angle-doubled Rayleigh vector length exceeds either 95% shuffle OR 0.18, a value derived from the population threshold (see below). Similarly, a TD-pattern cell was defined as meeting the following criteria: a). the cell’s peak firing rate ≥ 1 Hz; AND b). TD score exceeds the 95% shuffle; AND c). angle-quadrupled Rayleigh vector length exceeds either 95% shuffle or a population threshold at 0.15. Importantly, all criteria above were applied to both baseline trials in the 2-box and 4-box. The same criteria were also used in analyzing cell activity in 4-box darkness trials. For within-compartment classification, we further analyzed each cell’s firing in single compartments in the 1st baseline trials. In addition to meeting the overall criteria above, a between-compartment (BC) BD cell was defined as meeting these additional criteria: a). the Rayleigh vector length (without angle-doubling) in each single compartment exceeds 95% shuffle per cell or the mean Rayleigh vector length of single compartments exceeds 0.18 (population threshold); AND b). BD score in each single compartments did not exceed the shuffle control per cell. A within-compartment (WC) BD cell needs to meet these additional criteria: a). the mean angle-doubled Rayleigh vector length of single compartments exceeds 0.18; AND b). BD score in each single compartment exceeds 0.26 (population threshold). Similarly, for between-compartment (BC)TD cells: a). Rayleigh vector length (without angle-quadrupling) in single compartments exceeds 0.15 (population threshold) in at least three single compartments; AND b). TD score of single compartments did not exceed the shuffle control. For within-compartment (WC)TD cell: the TD score and angled-quadrupled Rayleigh vector length of single compartments exceeds 0.15 in at least three single compartments. In the 1-boxes (circular baselines), cells that were considered broad-unidirectional had their area under the correlation curve higher than its 95th percentile criterion of shuffle. Cells with their Rayleigh vector lengths higher than the 95% shuffle in all three successive cylinder trials were included for comparison analysis with HD cells.
‐ *Symmetry analyses:* To examine the directional pattern (onefold, twofold and fourfold) across environments and their sub-compartments, we calculated the cross-correlation (Pearson’s correlation coefficient) between two tuning curves in each angular bin in steps of 6°. For the 2-box, the tuning curve pairs were from lemon and vanilla sub-compartments of the baseline trials. For the 4-box, the tuning curve pairs were from any adjacent two sub-compartments (i.e., four pairs: vanilla 1 vs. lemon 1; lemon 1 vs. lemon 2; lemon 2 vs. vanilla 2; and vanilla 2 vs. vanilla 1) which were pooled together, and the final cross-correlation was averaged over these four comparisons. For 1-boxes, the tuning curve pairs were from two baseline trials. Periodicity would appear as a sinusoidal modulation of the circular cross-correlogram in a range from −180° to 180°, with the number of peaks reflecting the order of symmetry in the tuning curve. Computation for the shuffle control was performed in the same manner. Statistical tests were performed to compare the data and the shuffle.

##### 3. Egocentric spatial tuning analyses

We used the same set of analyses as described previously (*4, 13*). For each cell recorded in an environment, the animal’s allocentric position data points were used to get the following: movement direction (instantaneous derivative of successive position samples in the travelling path), the distance from a point to all environment boundaries visible by direct line-of-sight, and the angle from a point to all visible boundaries (in 3° increments). The spiking activity over the animal’s trajectories was color-coded by the animal’s movement directions.

*Egocentric rate map:* To construct the two-dimensional egocentric boundary rate maps, angular bins were referenced to the current movement direction so that 0° was to the animal’s front, 90° to its right, 180° to its back and −90° to its left. For every spike, its distance to the walls was computed within a range from zero to half of the length of the most distant possible boundary. The number of distance bins was 20, and the distance bin size and the maximum possible distance varied adaptively for each type of environments due to different environment dimensions (e.g., 40 cm for the circular arena, 50 cm for the square box, 60 cm for the 2-box and 30 cm for the 4-box). For a given 2D bin, the number of spikes within that spatial bin was divided by the dwell time. Rate maps were smoothed using a 2D Gaussian kernel (width = 5).

*Egocentric boundary tuning criteria:* We applied the same methods and criteria used previously (*4*). The mean resultant length was computed as the measure of egocentric boundary directionality. The mean resultant was calculated as:

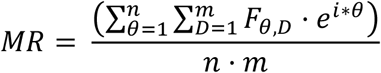

in which *θ* is the egocentric bearing to boundaries, *D* is the distance, *F*_*θ,D*_ is the firing rate in a given egocentric spatial bin, *n* is the number of orientation bins, *m* is the number of distance bins. Then MRL is calculated as the absolute value of MR. The mean resultant angle (MRA) - the preferred egocentric boundary orientation of a cell was calculated as:

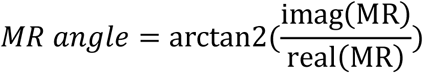

The preferred egocentric boundary distance was calculated based on MR angle by fitting a Weibull distribution to the firing rate along the MRA. The distance bin with the maximal firing rate was taken as the preferred boundary distance. For a cell to be a significant EBC: the MRL exceeds 99^th^ percentile of the shuffle, preferred distance shift less than 50% in the 1^st^ half vs. 2^nd^ half of a trial and MRA shift less than 45° in the 1^st^ and 2^nd^ half of a trial. We applied the above criteria to examine recorded directional cells and calculated the proportion that passed all three criteria in different environments.

##### 4. Basic electrophysiological features

The analyses were performed on the 1^st^ baseline trial of the multi-compartmented boxes for TD-pattern, BD-pattern and HD cells.

###### Waveform

The waveform width (peak-to-trough latency) and the amplitude were extracted for each cell. We collated the waveform features for all recorded cells and applied a K-means clustering method (MATLAB functions *kmeans)* to decide cluster centroid locations.

###### Burst index

The burstiness of neurons was measured by the spike-burst index, which was defined as the proportion of spiking activities (of all spikes) occurred within 6 ms of the inter-spike interval histograms (*14*).

###### Inter-spike interval

For each cell, the inter-spike interval of a spike train was analyzed within a time window of 50 ms. An epanechnikov kernel (bandwidth = 6) was fitted to the time interval distribution to get a smoothed density estimate. We calculated the latency between the points where the spiking probability first and last drops below half of the maximum, and then compared the population data between cell groups.

###### LFP phase and spiking activity

The LFP analysis methods were established and described in a previous study(*15*). To obtain a theta phase angle for each spike, the saved LFP signals were first bandpass filtered in the 6-12 Hz range (fourth-order Butterworth, MATLAB functions *butter* and *filtfilt*) before a Hilbert transform was applied to obtain the instantaneous phase angle (MATLAB function *hilber*t). For each cell, the instantaneous theta phase of every spike was calculated by linear interpolation of the instantaneous theta phase signal (MATLAB function *Phase*). The cell’s phase angles were binned between -π and π in 0.1 radian bins. The cell’s preferred theta phase was defined as the circular mean of these angles, and the strength of the modulation (phase-locking) was defined as the mean resultant vector length of these angles (MATLAB functions *circ_mean* and *circ_r* respectively (*10*)). For each cell, the number of spikes was normalized to its maximum in the baseline trial. At the population level, the average number of spikes per angular bin was calculated, and all cell preferred phases were collated.

#### Statistical analysis

We applied the circular statistic toolbox (*10*), and parametric tests and post hoc nonparametric tests were applied to compare population means and medians. In one-way ANOVA, Games-Howell post hoc procedure was used if the data violated the homogeneity of variance assumption. The statistical tests were performed in MATLAB (MathWorks, R2018) and SPSS packages (IBM, SPSS 25). Where appropriate the effect sizes for each test were stated clearly. All statistical tests are two-tailed (*p* value threshold at 0.05) unless stated otherwise. In all figures * = significant at the 0.05 level, ** = significant at the 0.01 level, *** = significant at the 0.001 level. For all box plots, red lines denote the sample mean, dotted lines denote the sample median, and boxes denote interquartile range.

#### Histology

At the end of the experiment, animals were anesthetized with isoflurane and injected with an overdose of sodium pentobarbital for euthanasia. Half of the animals underwent an electrolytic lesion by passing a small 15∼20 µA current through one to two electrical channels for 6∼12 seconds prior to perfusion. After a trans-cardiac perfusion, the brain was extracted and fixed in cold paraformaldehyde (4%). The brains were sliced using a freezing cryostat (Leica Biosystems, UK) under −21°C, and 35-40 μm coronal sections through caudal extent of the RSC were taken, stored in wells containing phosphate buffered saline (PBS), and wet-mounted on the slides. The slices best representing the electrode tracks were then imaged using an Olympus microscope (Olympus Keymed, UK). The deepest point of the electrode track was identified by referencing to the Paxinos and Watson rat brain atlas (Paxinos and Watson, 2006). The history of tetrode movements was used together with these points to calculate the tetrode depth for each recording session. The records were used to compare the localization of the recorded cells in dysgranular versus granular sub-regions in RSC.

## Supplementary figures and tables

Fig. S1. Experimental design.

Fig. S2. Directional tuning of BD-pattern cells in the 2-box experiment.

Fig. S3. Directional tuning of TD-pattern cells in the 4-box experiment.

Fig. S4. Within- and between-compartment patterns of TD cells in the 4-box.

Fig. S5. Maintained but decreased TD pattern in darkness.

Fig. S6. Comparison of multidirectional cell and HD cell activity.

Fig. S7. Multidirectional cells were not egocentrically tuned in any environment.

Fig. S9. Different distributions of multidirectional and HD cells in RSC sub-regions.

Fig. S10. Distinct electrophysiological properties of multidirectional cells vs. HD cells.

Table S1. Details of cell information in different environments.

Table S2. Statistical results comparing neural properties of multidirectional and HD cells.

**Fig. S1.**
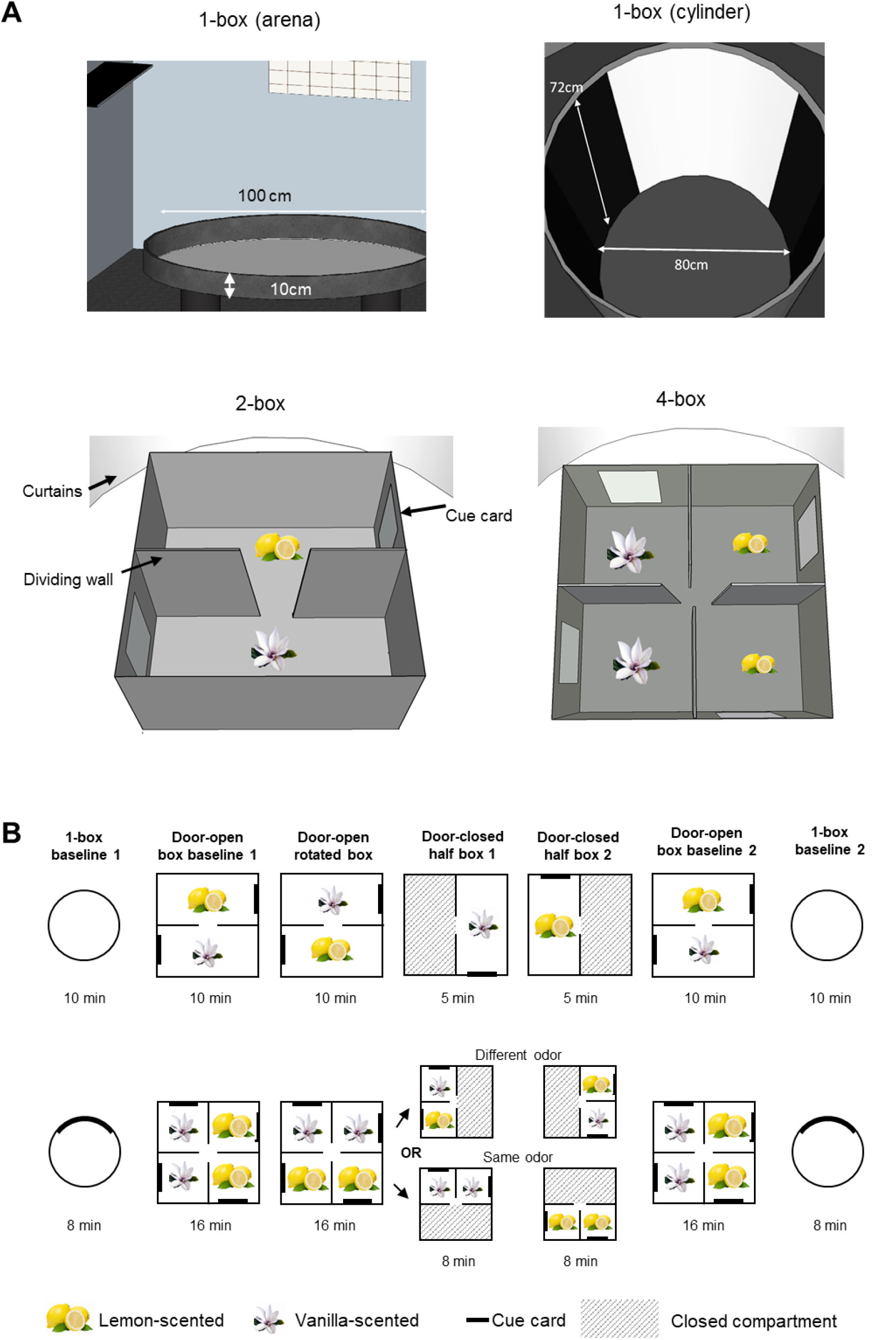
Experimental design. (A) Schematics of the environments. (B) Recording protocols for the 20box experiment (top) and 4-box experiment (bottom).

**Fig. S2.**
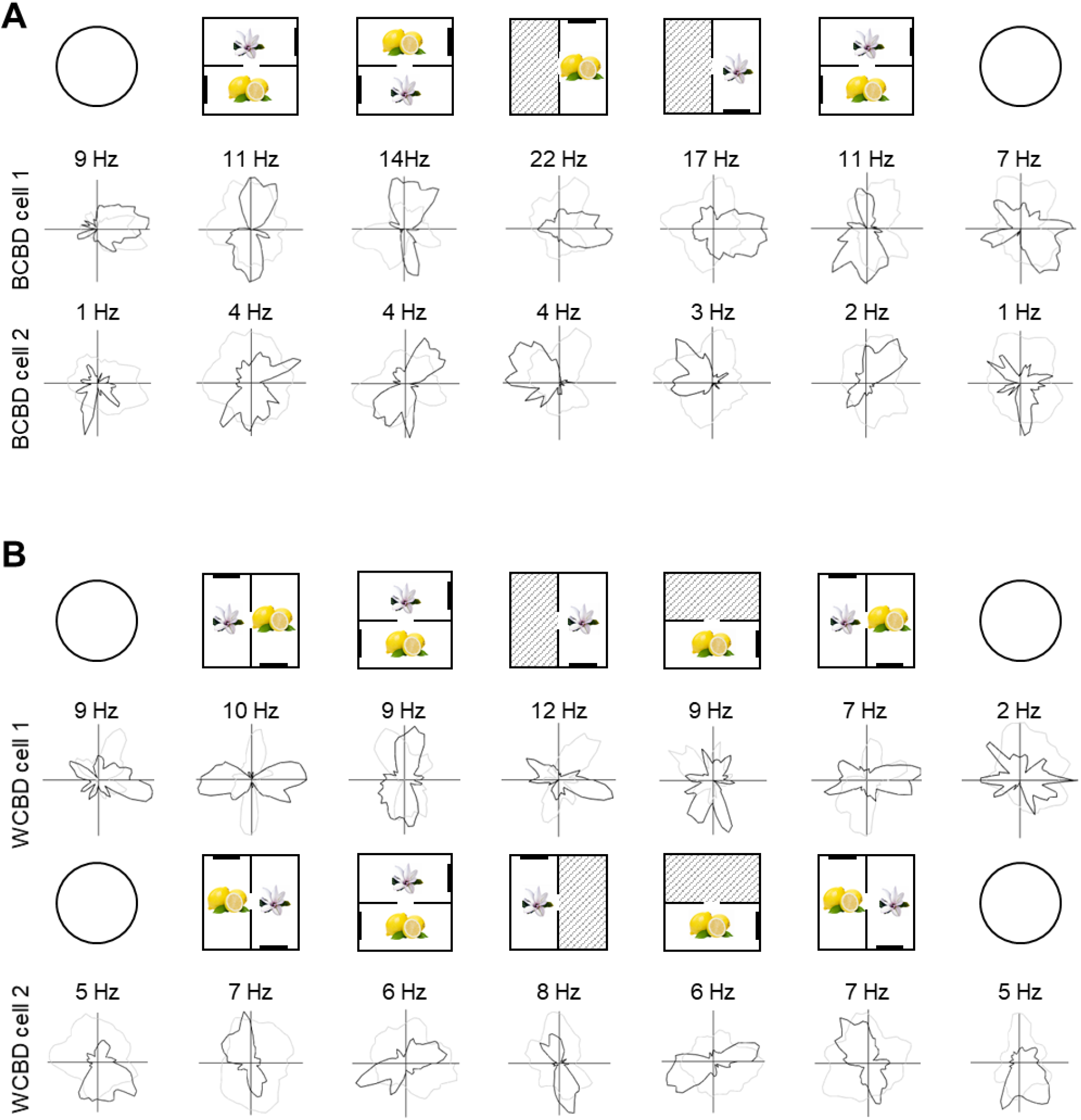
Directional tuning of BD-pattern cells in the 2-box experiment. Four example BD-pattern cells recorded from different animals in the 2-box experiment, the BCBD cells are shown in (A) and WCBD cells in (B). Polar plots show the standard seven-trial recording sessions. The 1-box here was the open arena. Note that the BD-pattern cell firing patterns were miscellaneous in the 1-box

**Fig. S3.**
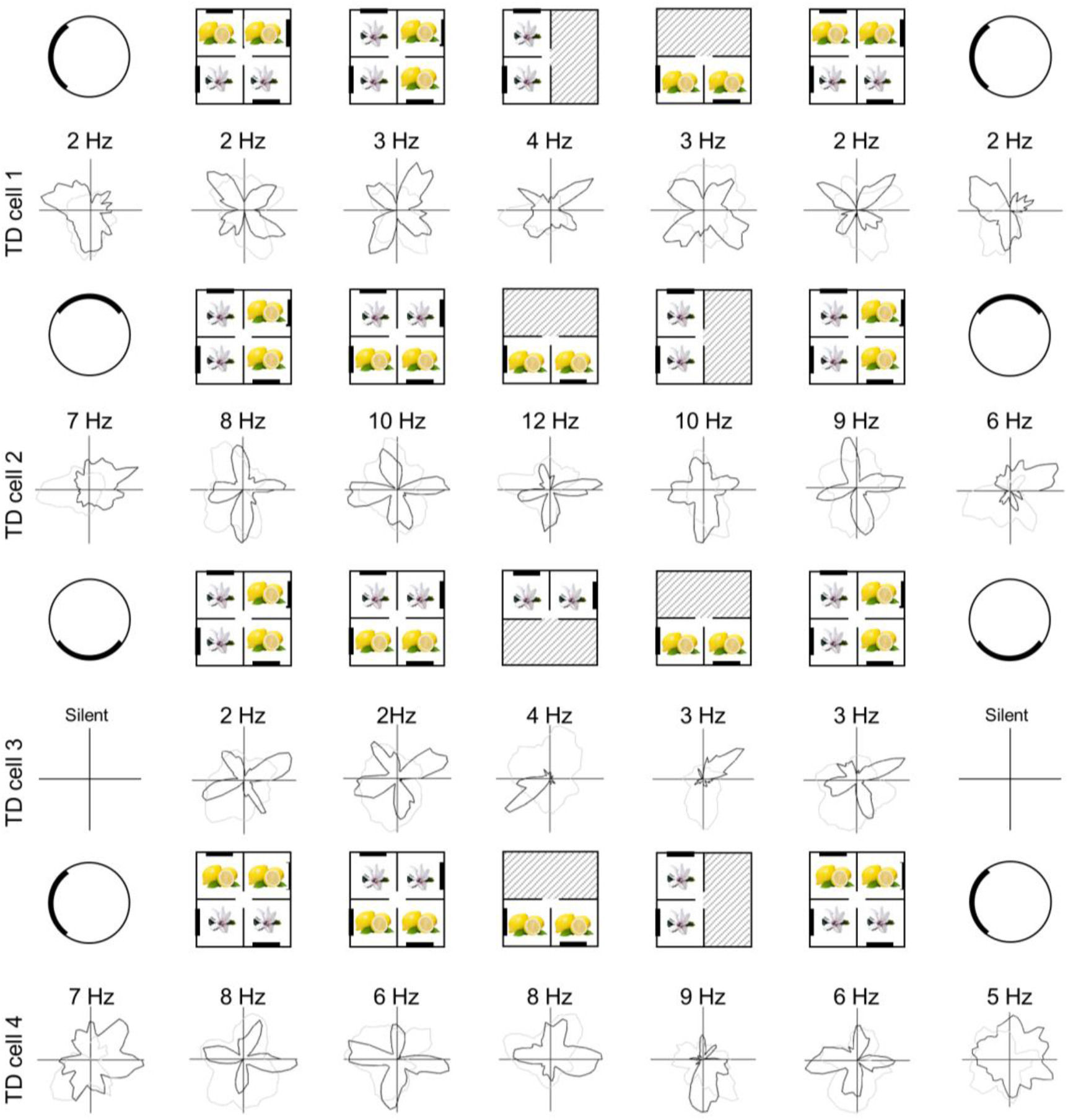
Directional tuning of TD-pattern cells in the 4-box experiment. Four example TD-pattern cells are shown, recorded from different animals in the 4-box experiment. Polar plots show the standard seven-trial recording sessions. Note that directional firing patterns are miscellaneous in the 1-box baseline trials, and one cell did not fire at all in these environments.

**Fig. S4.**
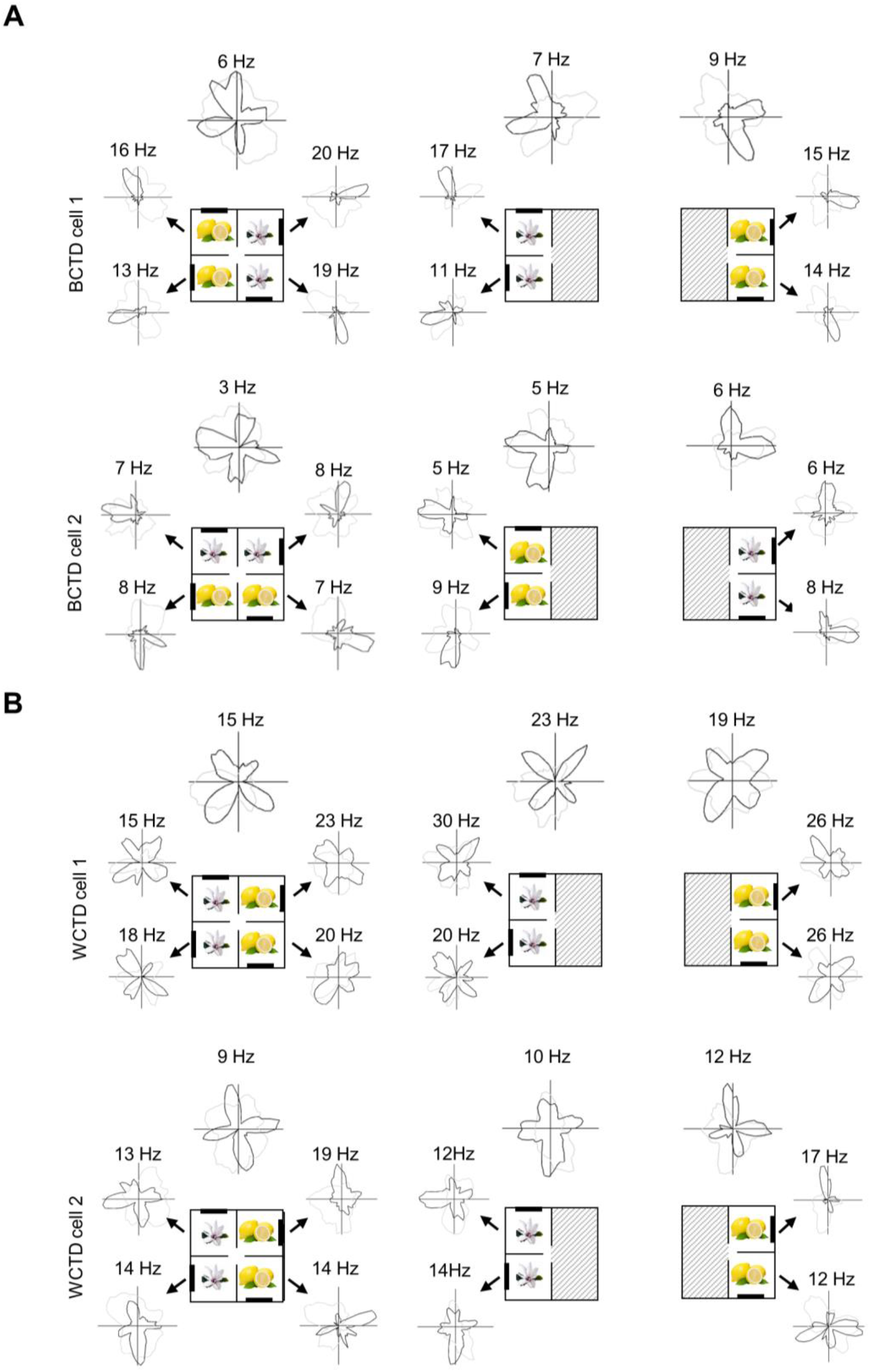
Within- and between-compartment patterns of TD cells in the 4-box. When firing patterns of TD cells were examined within individual compartments, two patterns emerged as with the BD cells. Some cells showed only a singlefold tuning in the single compartments (BCTD cell, two examples shown in A) and some maintained a fourfold pattern (WCTD cell, two examples shown in B).

**Fig. S5.**
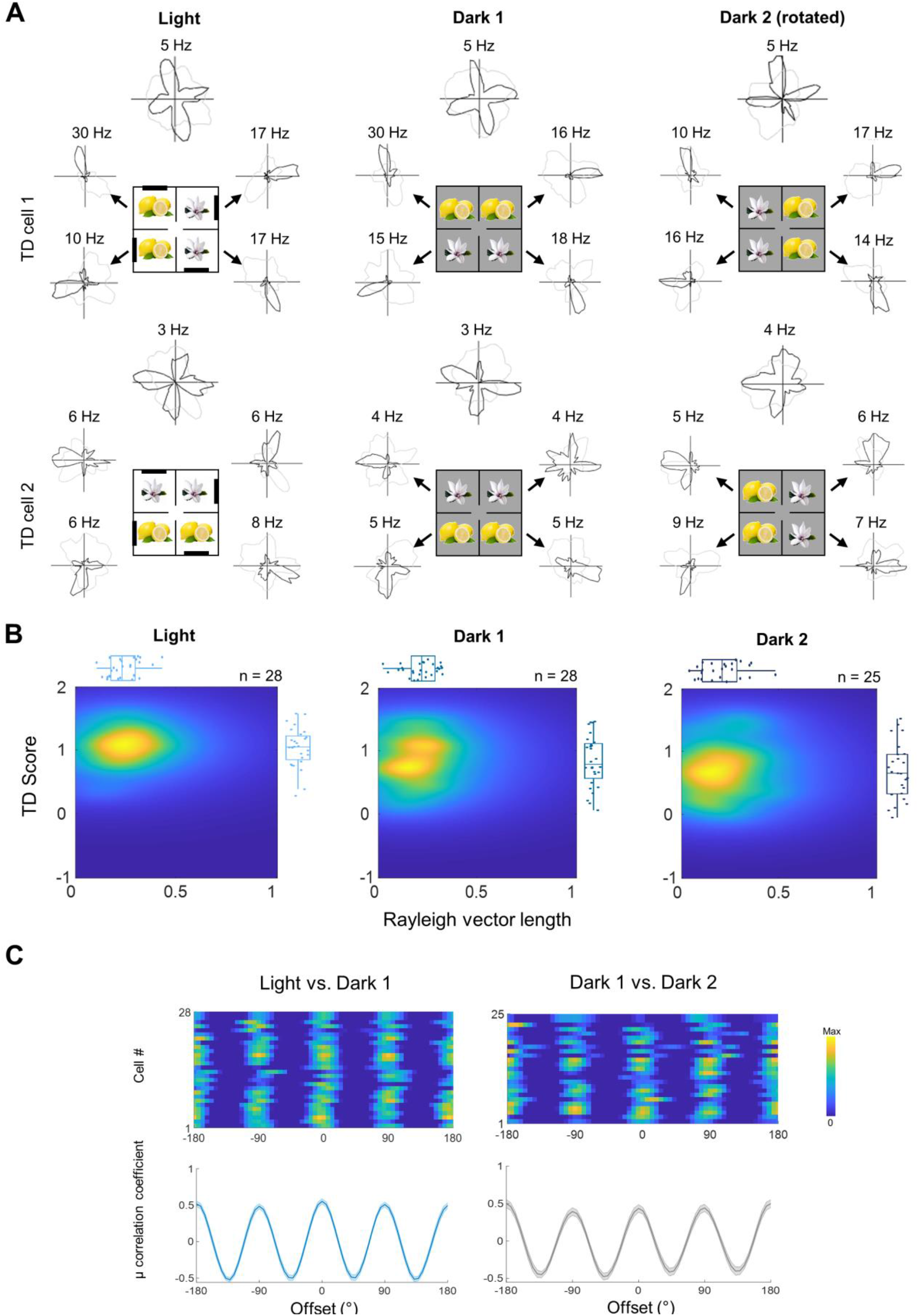
Maintained but decreased TD pattern in darkness. (A) Example of two TD-pattern cells that maintained the four-way firing pattern in two darkness trials following the 4-box baseline. Note that the animal was removed from the 4-box, mildly disorientated and reintroduced to the apparatus before the dark trials started. (B) Density and box plots showing distribution of TD scores as a function of Rayleigh vector length in light (1^st^ box baseline) and two dark trials – significantly different in TD scores (repeated-measures ANOVA *F*(2,78) = 5.26, *p* = 0.007) and Rayleigh vector length within the group (*F*(2,78) = 3.36, *p* = 0.04). Between-group comparisons of TD scores: significant for light vs. dark 1 (t-test t(27) = 2.51, *p* = 0.018) and dark 1 vs. dark2 (t(24) = 2.54, *p* = 0.018). Between-group comparisons of Rayleigh vector length: significant for light vs. dark 1 (t(27) = 4.35, *p* < .001) but non-significant between two dark trials (t(24) = −0.52, *p* = 0.61). (C) Cross-correlograms of tuning curves the mean correlation coefficients of light vs. dark 1 (left, blue) and two dark trials (right, grey), and they were not significantly different: KStest, D = 0.165, *p* = 0.142.

**Fig. S6.**
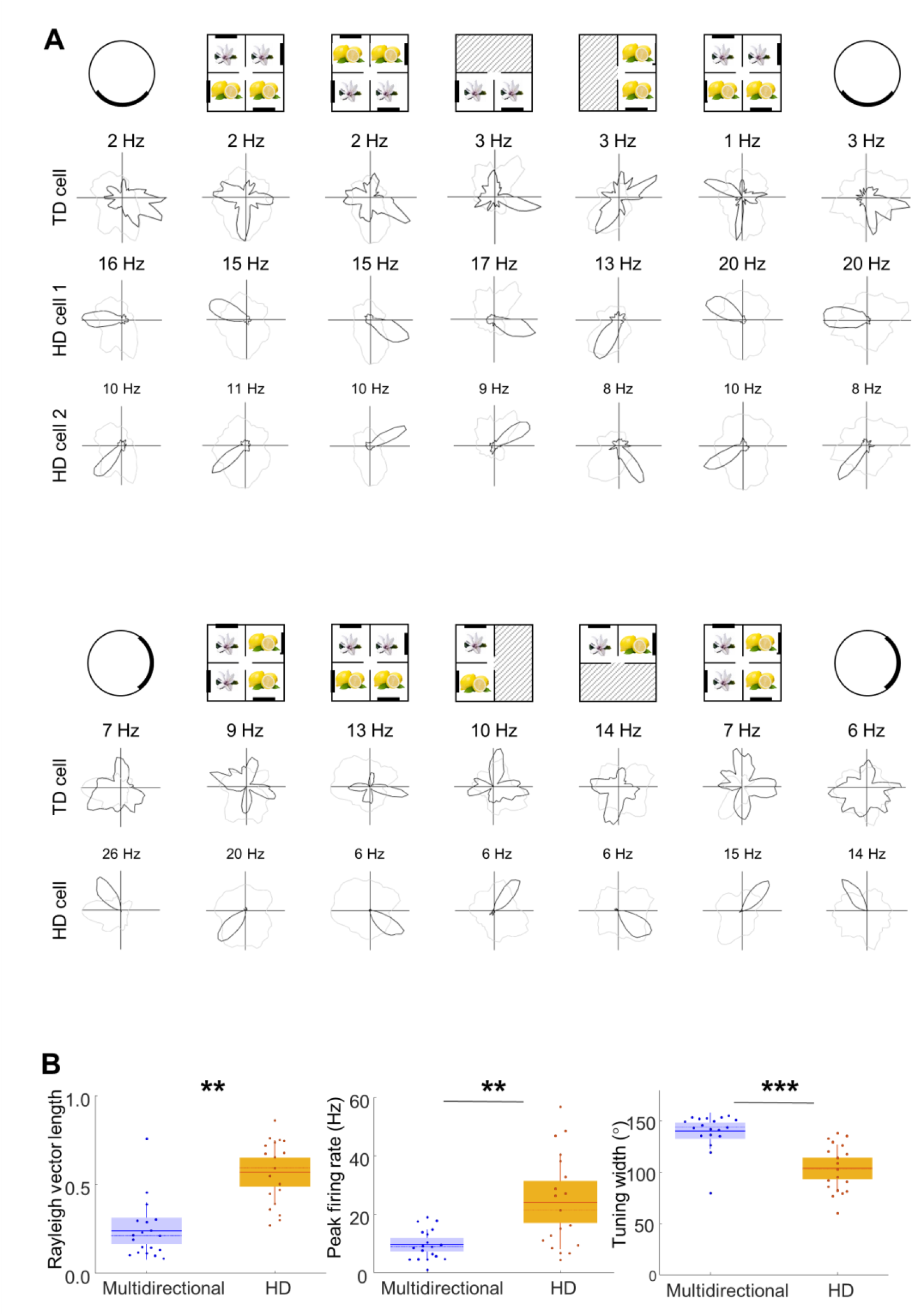
Comparison of multidirectional cell and HD cell activity. (A) Examples of one TD-pattern co-recorded with two HD cells in one session (upper) and another TD-pattern co-recorded with an HD cell in a different recording session (bottom). Simultaneous recordings of the HD cell and the TD-pattern cell reveal dissociation in direction encoding of the two groups. (B) Comparison of multidirectional and HD cell activity in 1-box trials: box plots show directional tuning characteristics of 19 multidirectonal cells (15 TD-pattern and 4 BD pattern, violet) and HD cells (orange). Rayleigh vector length (left): multidirectional, median = 0.21; HD, median = 0.59; *Wilcoxon rank-sum* Z = −4.34, *p* <.001; Peak firing rate (middle): multidirectional, median = 8.9Hz; HD, median = 21.4Hz; Z = −2.95, *p* = 0.003), and tuning width (right): multidirectional, median = 144°; HD, median = 103°; Z = 4.38, *p* <.001).

**Fig. S7.**
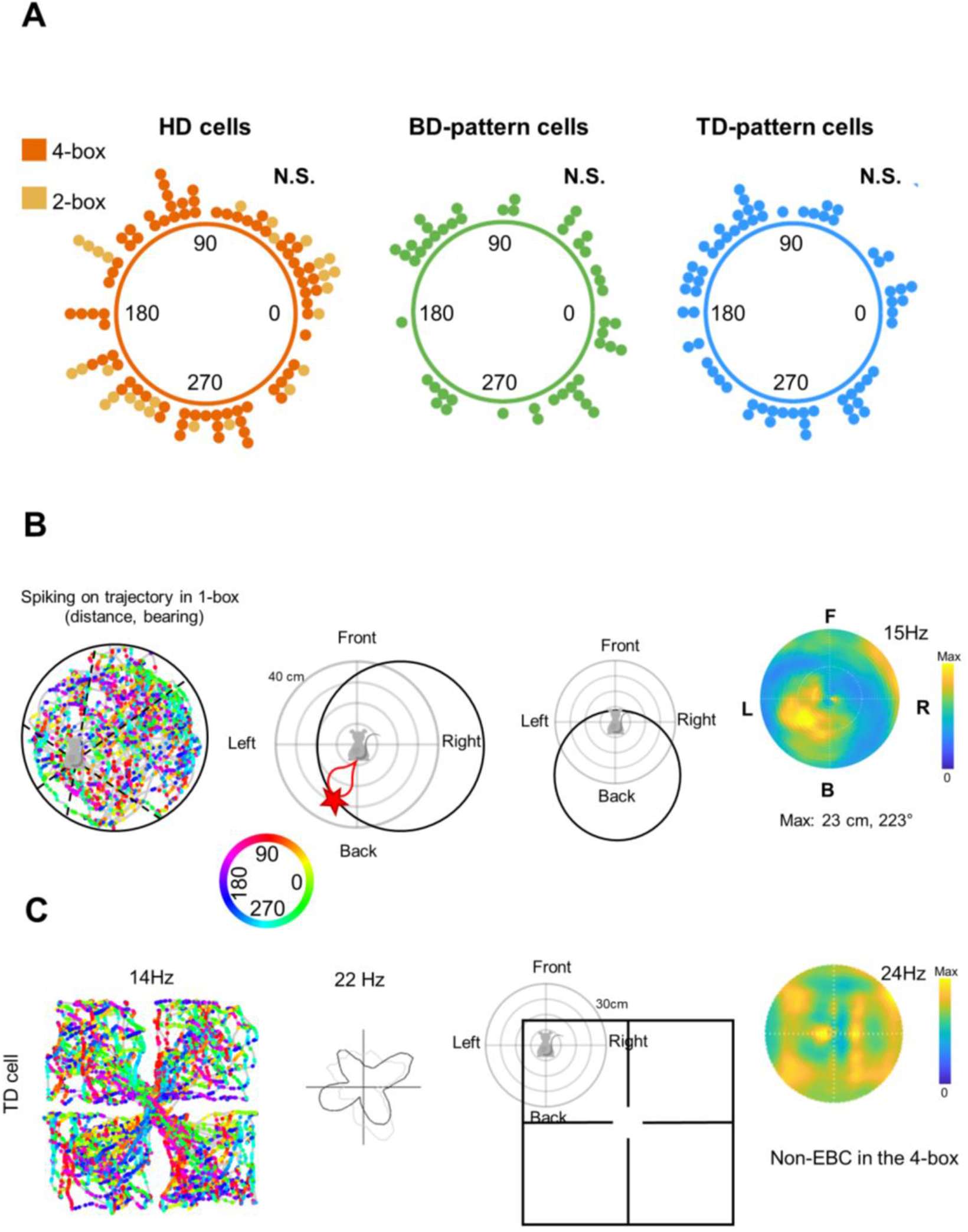
Multidirectional cells were not egocentrically tuned in any environment. (A) Allocentric HD tuning of retrosplenial cells was homogeneously distributed in the multifold environments. Circular distribution of preferred directions of unidirectional HD cells (orange), (the largest peak of BD-pattern (green) and TD-pattern (blue) in the multicompartmented environments. Note that the preferred orientations of the multidirectional cells and HD cells were distributed homogenously across the whole environment, Rayleigh test: TD: z = 0.76, p = 0.47; BD: z = 0.63, p = 0.53; HD: z = 0.04, p = 0.96).

**Fig. S8.**
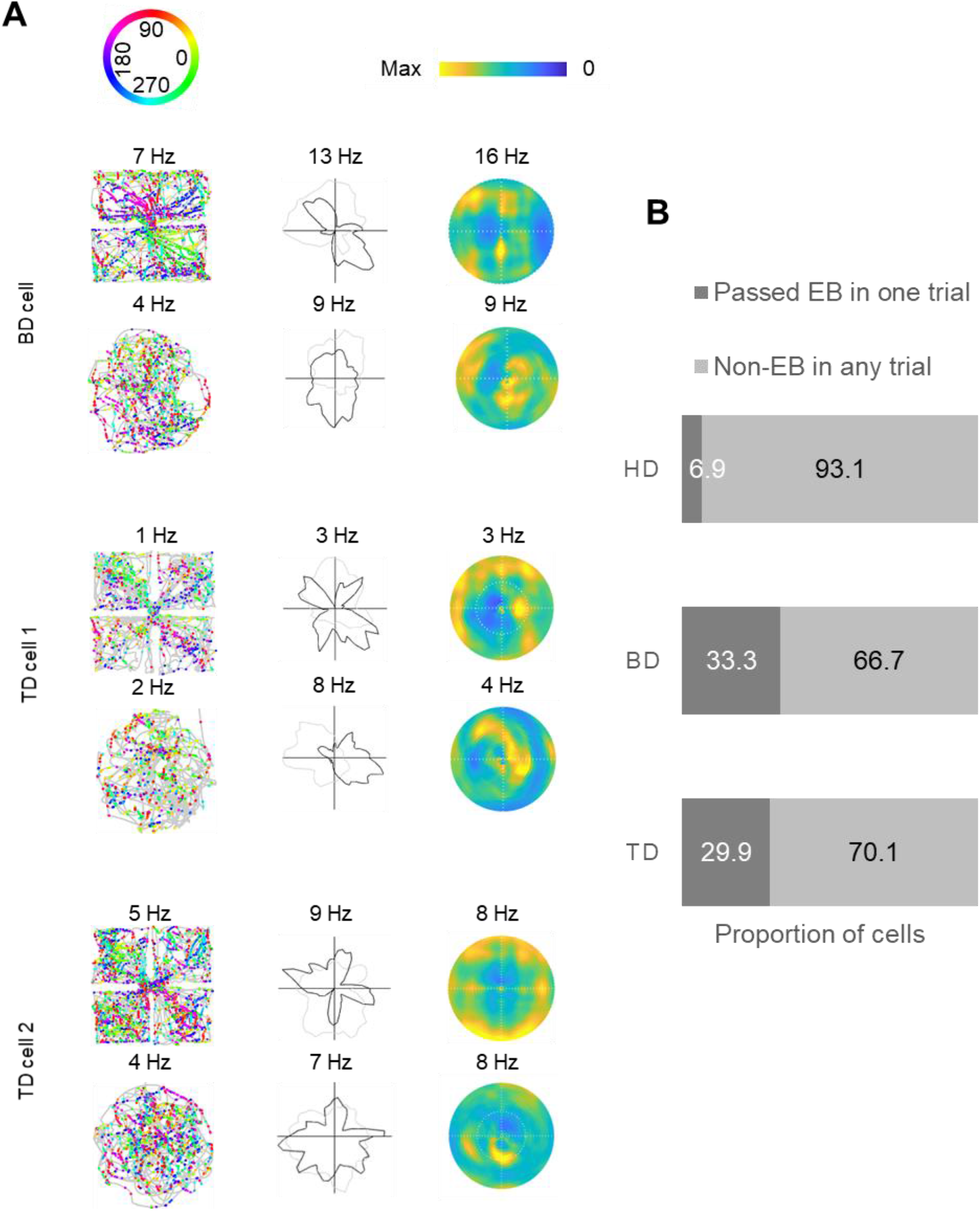
(A) Example of one BD pattern cell (top) and two TD-pattern cells (middle and bottom) and one BD-pattern cells, recorded in a multi-compartment or single arena. Left: spike plots, color-coded by allocentric facing direction. Egocentric boundary (EB) heatplots (right) reveal homogeneous firing with dispersed fields and no clear hot spot. (B) Proportion of directional cells (%) assessed as passing (dark gray) or failing (pale gray) the EB criteria in two baseline trials in the 1-box and multifold boxes. The majority of directional cells did not pass the EB criteria in any trial, and a significantly higher amount of HD cells than multidirectional cells failed to meet the threshold (Chi-Squared test: TD vs HD: χ2(1, N = 169) = 15.92, *p* <.001; BD vs HD: χ2(1, N = 150) = 17.62, *p* <.001; TD vs BD: χ2(1, N = 115) = 0.16, *p* = 0.691).

**Fig. S9.**
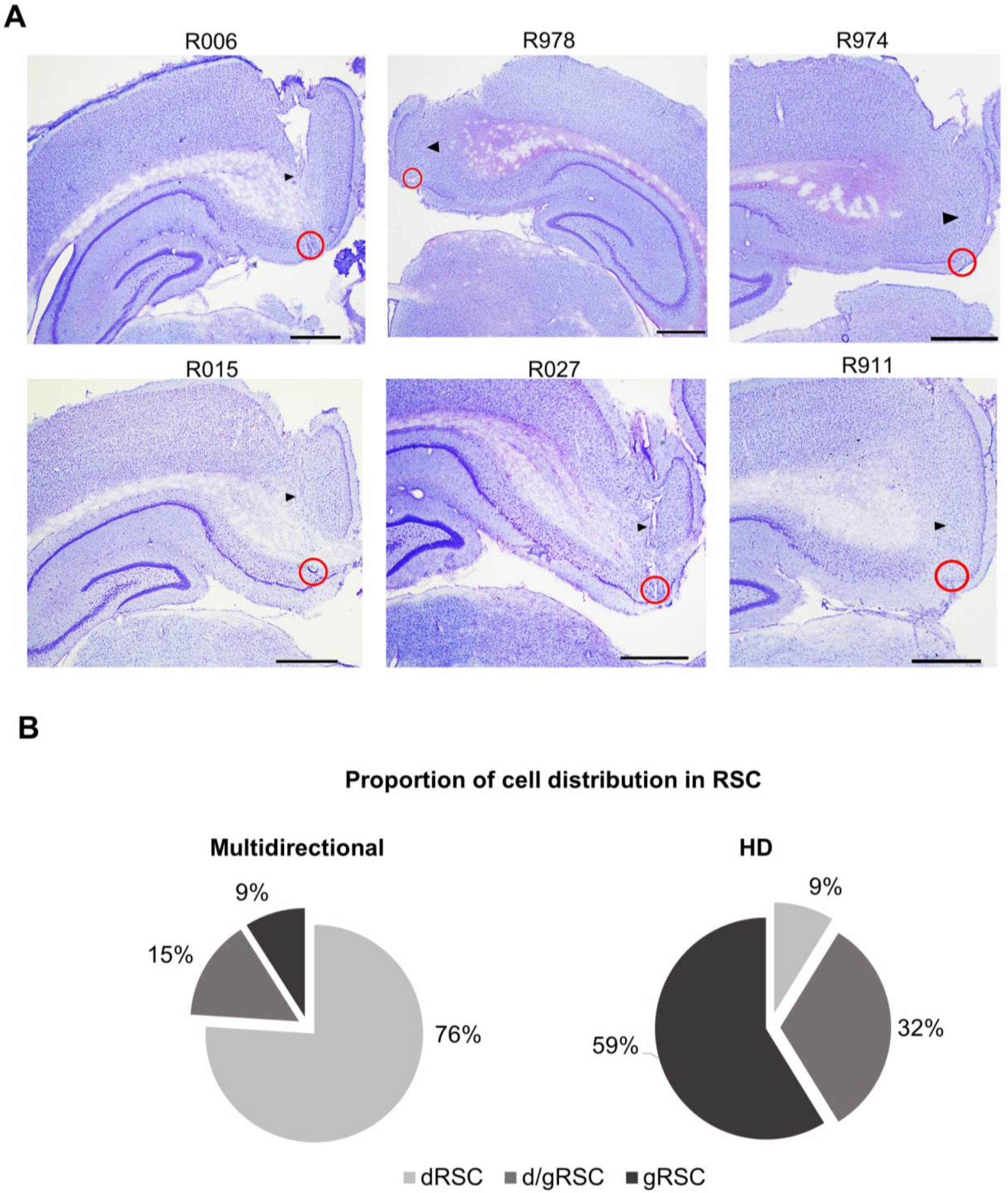
Different distributions of multidirectional and HD cells in RSC sub-regions. (A) Example histological slices of six animals implanted with 8-tetrode (R006, R978, R015, and R027) or 4-tetrode bundles (R974, R911). Animals of these representative sections received a small electrolytic lesion under anesthesia prior to perfusion, showing visible brain tissue damage at the tip of electrodes, indicated by red circles. Black arrow points to the trajectory of the electrode bundle. Scale bar denotes 1mm. The median recording coordinates were: AP: −5.5, ML: ±0.83 of all animals (n=18). (B) Proportion of the directional cells shown in pie charts (left: multidirectional cells, n = 115; right: HD cells, n = 102) recorded in each RSC sub-region (light grey: dysgranular RSC; dark grey: intersection between dysgranular and granular RSC; space grey: granular RSC, Chi-Squared test: χ^2^ (2, N=217) = 101.89, *p* <.001).

**Fig. S10.**
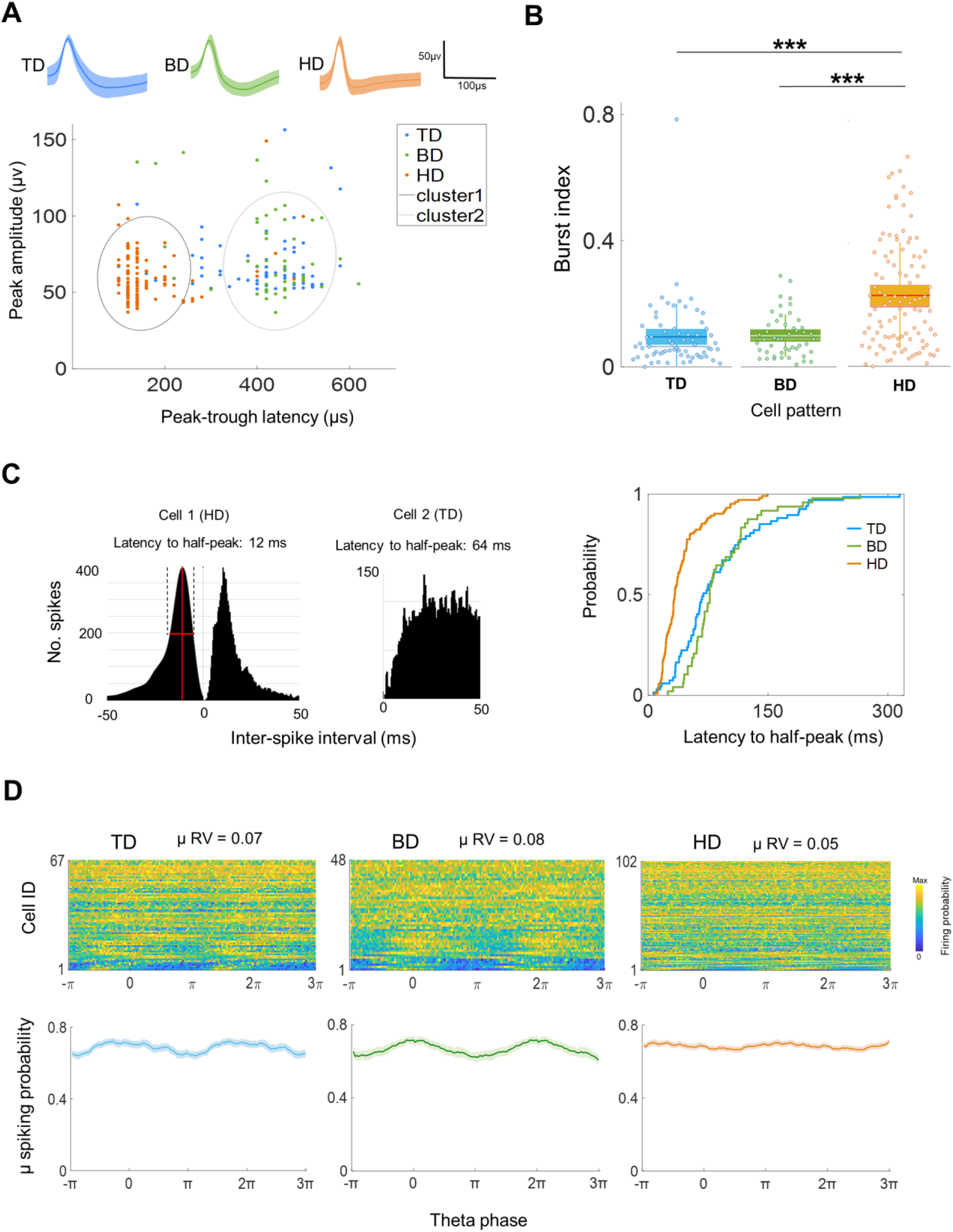
Electrophysiological properties of multidirectional cells are distinguishable from HD cells. (A) Top: Example waveforms of a TD-pattern (blue), BD-pattern (green) and HD cell (orange) of the channel with the highest amplitude (Mean± s.e.m); numbers denote of waveform peak amplitude and peak-trough latency. Bottom: scatter plots show the multidirectional cell cluster (cluster 1) distinct from HD cells (cluster 2), K-Means clustering, best total sum of distance between two clusters = 11946.3. The distributions were plotted as ellipses specified by population mean and covariance. The two separated distributions were significantly different in their peak-trough latency (t(215) = 38.79, *p* < .001) and peak amplitude (t(215) = 2.88, *p* =0.0044). (B) Box plots of burst index show significantly different distributions between TD and HD, BD and HD cells, but not between TD and BD (solid line: mean, dotted line: median). (C) Left: Example of an HD cell (left) and TD-pattern cell (right), showing distinct spiking distribution in inter-spike intervals analysis. Right: cumulative probability distributions of latency to half-peak (FWHM) for TD (blue) and BD (green) cells were similar whereas different from HD cells (orange). (D) Top: Heatmaps show spiking activity as a function of theta phase for three patterns. Cell ID was sorted by their Rayleigh vector lengths (maximum at the bottom), representing the strength of phase-locking. The number denotes the population mean strength of phase-locking. Bottom: Normalized spiking activity (mean±SEM) over two theta phases. The relationship between theta and cell spiking was not significantly different between the BD vs. TD groups or TD vs. HD groups, but it was slightly different between BD vs. HD groups; See statistical summary in Table S2.

**Table S1:**
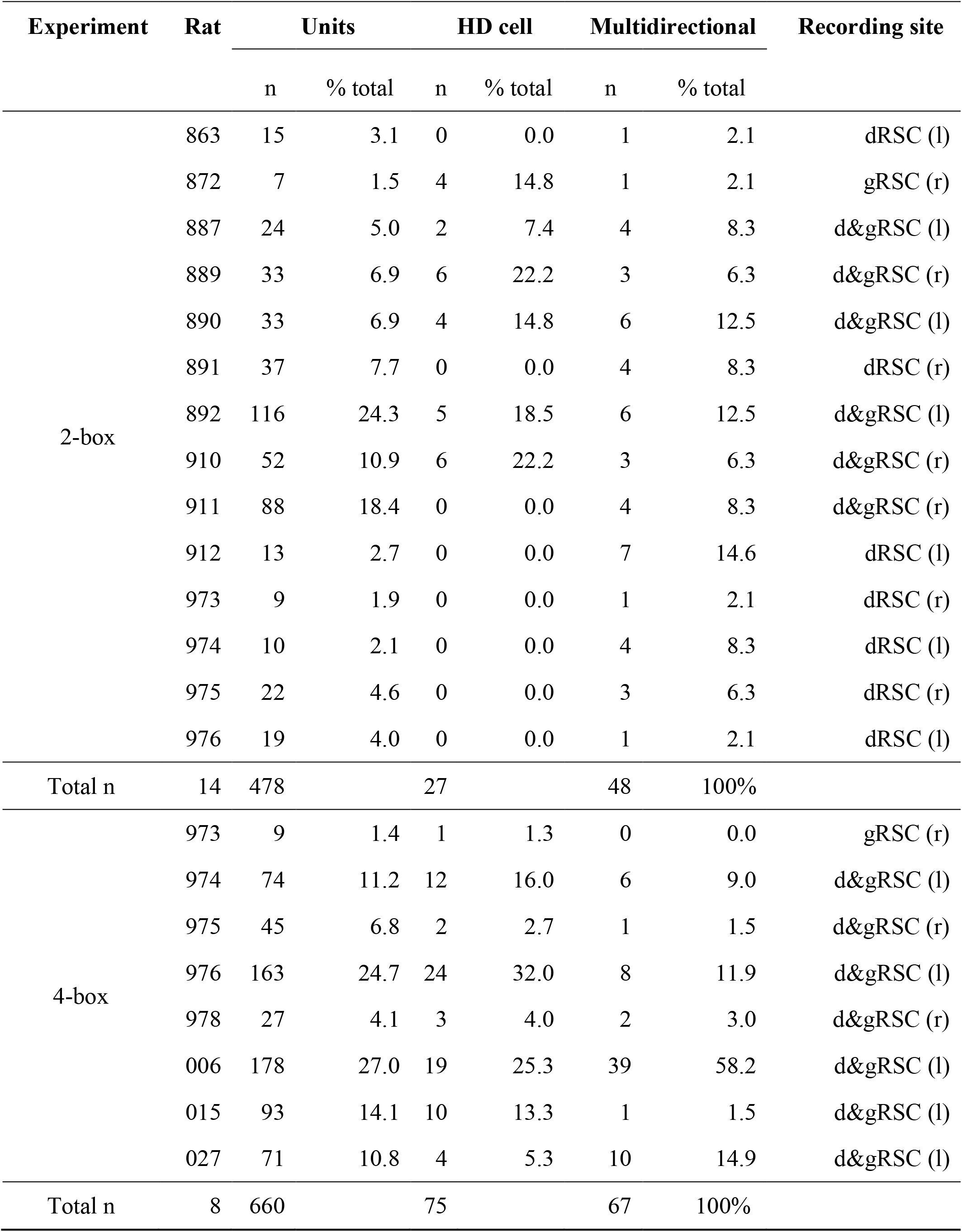
Data summary.

**Table S2:** Electrophysiological firing properties of directional cells.

| Neural properties | TD pattern | BD pattern | HD cells | Welch's ANOVA* | Post-hoc comparisons |
| --- | --- | --- | --- | --- | --- |
| Peak amplitude (μV) | 68.51±2.35 | 74.69±3.94 | 60.40±1.64 | $F(2, 101.87) = 7.84$,<br>$p = 0.001$ | TD vs BD:<br>$p = 0.374$;<br>TD vs HD:<br>$p = 0.015$;<br>BD vs HD:<br>$p = 0.004$. |
| Peak-trough latency (μs) | 402.39±13.72 | 405.83±17.51 | 162.75±7.99 | $F(2, 100.98) = 159.73$, $p < .001$ | TD vs BD:<br>$p = 0.987$;<br>TD vs HD:<br>$p < .001$;<br>BD vs HD:<br>$p < .001$. |
| Burst index | 0.10±0.01 | 0.10±0.01 | 0.23±0.02 | $F(2, 140.61) = 24.81$, $p < .001$ | TD vs BD:<br>$p = 0.926$;<br>TD vs HD:<br>$p < .001$;<br>BD vs HD:<br>$p < .001$. |
| Latency to ISI half-peak (ms) | 88.17±7.28 | 89.67±6.62 | 42.12±2.91 | $F(2, 94.87) = 33.64$, $p < .001$ | TD vs BD:<br>$p = 0.987$;<br>TD vs HD:<br>$p < .001$;<br>BD vs HD:<br>$p < .001$. |
| Theta modulation strength | 0.07±0.01 | 0.08±0.01 | 0.05±0.005 | $F(2, 108.37) = 2.98$,<br>$p = 0.055$ | TD vs BD:<br>$p = 0.523$;<br>TD vs HD:<br>$p = 0.252$;<br>BD vs HD:<br>$p = 0.07$. |

\* The assumption of homogeneity of variances was violated, and so Welch's ANOVA was used as an alternative to one-way ANOVA. Games-Howell test was used to compare all possible combinations to detect group differences.

## Notes

### Summary of Updates

The authors have added the supplementary materials.

